# Spatial transcriptomics reveals molecular dysfunction associated with Lewy pathology

**DOI:** 10.1101/2023.05.17.541144

**Authors:** Thomas Goralski, Lindsay Meyerdirk, Libby Breton, Laura Brasseur, Kevin Kurgat, Daniella DeWeerd, Lisa Turner, Katelyn Becker, Marie Adams, Daniel Newhouse, Michael X. Henderson

## Abstract

Lewy pathology composed of α-synuclein is the key pathological hallmark of Parkinson’s disease (PD), found both in dopaminergic neurons that control motor function, and throughout cortical regions that control cognitive function. Recent work has investigated which dopaminergic neurons are most susceptible to death, but little is known about which neurons are vulnerable to developing Lewy pathology and what molecular changes an aggregate induces. In the current study, we use spatial transcriptomics to selectively capture whole transcriptome signatures from cortical neurons with Lewy pathology compared to those without pathology in the same brains. We find, both in PD and in a mouse model of PD, that there are specific classes of excitatory neurons that are vulnerable to developing Lewy pathology in the cortex. Further, we identify conserved gene expression changes in aggregate-bearing neurons that we designate the Lewy-associated molecular dysfunction from aggregates (LAMDA) signature. This gene signature indicates that neurons with aggregates downregulate synaptic, mitochondrial, ubiquitin-proteasome, endo-lysosomal, and cytoskeletal genes and upregulate DNA repair and complement/cytokine genes. However, beyond DNA repair gene upregulation, we find that neurons also activate apoptotic pathways, suggesting that if DNA repair fails, neurons undergo programmed cell death. Our results identify neurons vulnerable to Lewy pathology in the PD cortex and identify a conserved signature of molecular dysfunction in both mice and humans.

## INTRODUCTION

Parkinson’s disease (PD) is diagnosed post-mortem by loss of dopaminergic neurons in the substantia nigra pars compacta (SNc) and the presence of α-synuclein Lewy pathology throughout the brain^1^. Yet, it is unclear what is happening in neurons with Lewy bodies (LBs), and some have argued that these large cytoplasmic inclusions may even be protective (reviewed here^2^). Further, outside of the SNc, there is minimal knowledge about which neurons are susceptible to developing Lewy pathology. The current study aimed to discover which neurons bear α-synuclein inclusions and what molecular processes are changed in these neurons using spatial transcriptomic and complementary approaches in a mouse model of α-synucleinopathy and in PD brain.

PD is diagnosed clinically by the presence of motor dysfunction^1^, however, PD patients experience many other symptoms. One of the most devastating symptoms of PD is the progression of over 80 percent of patients to dementia^3^, a diagnosis associated most strongly with progression of Lewy pathology to cortical regions^4–8^. Many brainstem nuclei sustain neuron loss in PD, and vulnerability factors have been identified that unite several of these nuclei—autonomous pacemaking activity, hyperbranching axons, and synthesis of catecholaminergic neurotransmitters that may promote mitochondrial dysfunction^9, 10^. Yet, even within these highly susceptible types of neurons, there is a selective vulnerability. Early morphological studies found that the ventral tier of the SNc is particularly vulnerable to neuron loss in PD^11, 12^. Advances in RNA sequencing have shown that *SOX-AGTR1*-positive neurons are selectively lost in the SNc^13^ and there is a dysfunctional transcriptomic signature associated with SNc neurons in PD^13–15^. Yet, these scRNAseq studies did not assess molecular processes in neurons bearing Lewy pathology.

Staging studies have shown that Lewy pathology extends past the brainstem to neocortical regions in later stages, especially cingulate and prefrontal cortex^16^. Within those regions, there is a sparse distribution of affected neurons mostly in deep layers^16^, suggesting that subpopulations of neurons in those layers are particularly affected. These appear to be cortico-cortical projection neurons because both short-axoned projection neurons, inhibitory interneurons, and large myelinated cortico-spinal neurons do not develop pathology^17^. This has suggested that, as with catecholaminergic neurons, a long axon is a potential source of vulnerability, although myelination of corticospinal neurons may be protective^17^.

We hypothesized that there are select populations of neurons vulnerable to developing Lewy pathology in PD and that those neurons would show disruption of cellular processes preceding their loss. To separate α-synuclein inclusion-bearing from resilient neurons, we used immunostaining and spatial transcriptomics to collect whole-transcriptome molecular signatures from those two populations in a mouse model of α-synucleinopathy and PD brains. We were able to clearly identify transcriptomic changes by cortical layer and by inclusion status. Layer 5 intratelencephalic (IT) and layer 6b neurons were particularly vulnerable, while inhibitory and pyramidal tract (PT) neurons were resilient. Further, inclusion-bearing neurons had reduced synaptic, mitochondrial, proteasomal, endo-lysosomal, and cytoskeletal gene expression, while genes associated with DNA damage repair, apoptosis, and complement cascade were elevated. In PD brain, we showed similar ability to detect cortical layer-selective genes and found a similar distribution of Lewy pathology, predominantly in layer 5 within IT neurons, while inhibitory and pyramidal neurons were spared. Molecular changes in LB-bearing neurons showed remarkable similarity to those seen in the α-synucleinopathy mouse model, with over 600 conserved gene expression changes. Again, genes associated with the synapse, mitochondria, proteasome, endo-lysosome and cytoskeleton were down-regulated, while DNA damage repair, apoptosis, and complement cascade genes were upregulated. Together, this study indicates that there is a selectively vulnerable population of neurons in PD cortex, and α-synuclein inclusions are associated with a strong and conserved stress response in neurons. Future studies may take advantage of this knowledge to target specific classes of neurons and molecular pathways for therapeutic effect.

## RESULTS

### Collection of whole transcriptome signatures of neurons with or without pathology

Single-cell transcriptomics has enabled the examination of gene signatures associated with PD^13–15, 18, 19^. However, none of these studies have examined whether or not the isolated neurons bear α-synuclein inclusions. Isolating inclusions is difficult using such techniques due to the necessary dissociation of cells, and the capture of only nuclei for most post-mortem human brain studies. We therefore employed a recentl-developed Whole Transcriptome Atlas capture base on *in situ* hybridization technology using the GeoMx digital spatial profiler (DSP)^20^.

We first assessed the pathology distribution in mice injected with α-synuclein pre-formed fibrils (PFFs) in the dorsal striatum (**Fig. 1A**). Mice were aged 3 months post-injection and developed a reproducible distribution of α-synuclein inclusions in frontal cortical regions—anterior cingulate (ACA), secondary motor (MOs), and primary motor (MOp, **Fig. 1B**) cortex. Further, inclusion burden was most densely distributed in cortical layer 5 of these regions. In order to segment neurons with and without pathology, sections were stained with neuron marker NeuN, astrocyte marker glial fibrillary acidic protein (GFAP), and pathology marker pS129 α-synuclein (pSyn, **Fig. 1C, Supplementary Fig. 1**). To accurately annotate cortical layers, brain section images were downloaded from the GeoMx instrument, registered to the Allen Brain Atlas CCFv3 using a modified version of the QUINT workflow^21, 22^, and registrations were overlaid on the slide image in the GeoMx instrument (**Fig. 1C**). Regions of interest were drawn within Allen Brain Atlas regions. Segmentations were developed to identify two classes of neurons, those that were NeuN+/pSyn-and those that were NeuN+/pSyn+ (**Fig. 1C**). These two classes are referred to as NeuN (pathology free) and pSyn (pathology-bearing) for the remainder of the manuscript.

**Figure 1.**
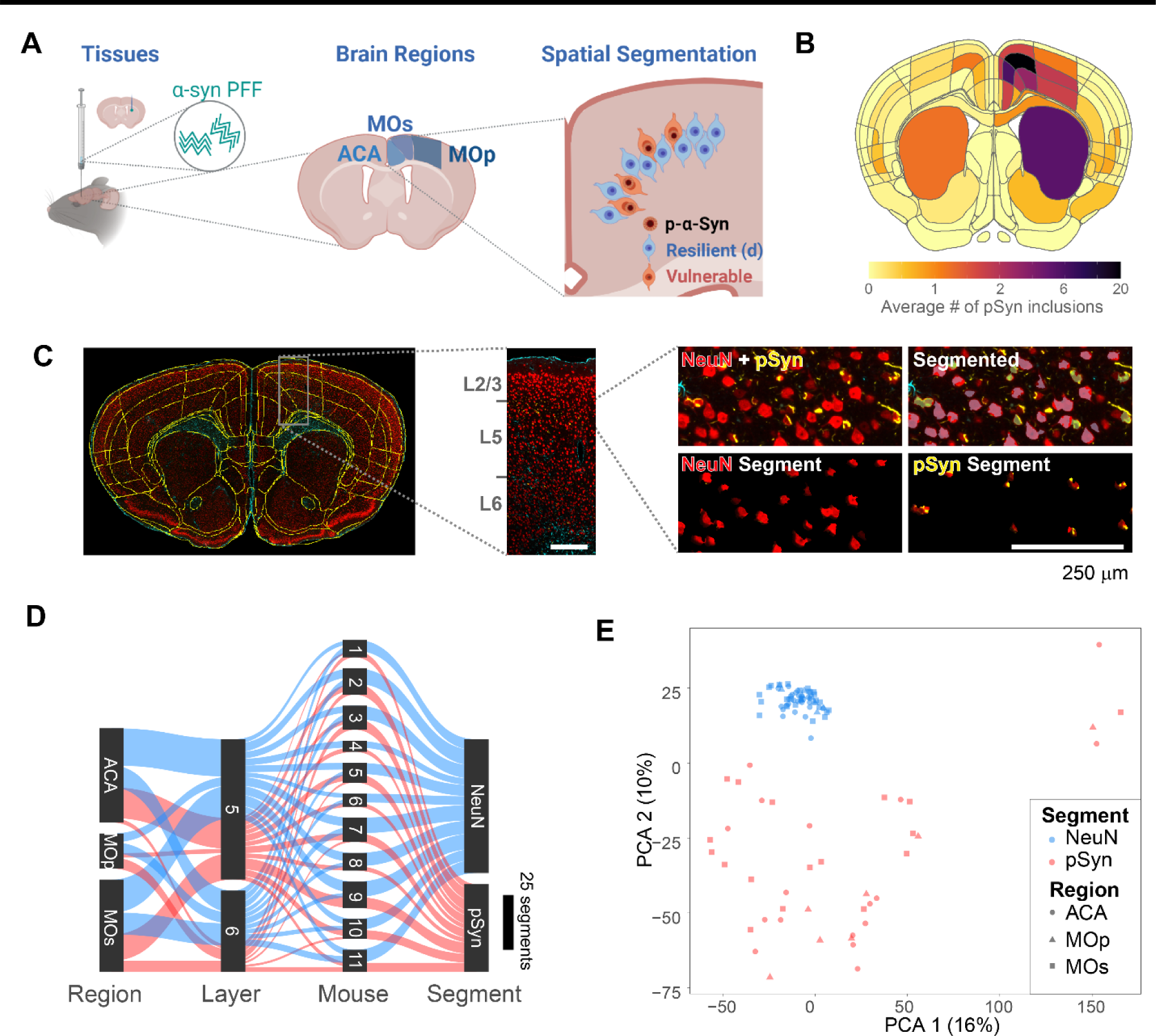
Collection of transcriptome signatures of neurons with or without pathology with GeoMx. (**A**) Experimental schematic. Mice were injected in the dorsal caudoputamen with α-synuclein PFFs. After 3 months, the brains of mice were removed and brain sections were prepared for spatial transcriptomic analysis, with a focus on frontal cortical regions—anterior cingulate (ACA), secondary motor (MOs), and primary motor (MOp). Within each region vulnerable or resilient neurons were identified by the presence or absence of α-synuclein pathology and RNA from those regions were selectively identified. (**B**) Quantification of the average number of pSyn inclusions per region from 12 mice injected with α-synuclein PFFs. ACA, MOs, and MOp had the highest pathology burden, especially cortical layer 5. (**C**) A representative brain section stained for NeuN (red), GFAP (cyan), and pSyn (yellow) was registered to the Allen Brain Atlas CCFv3. A zoomed view of MOs/ACA is shown denoting layers of the cortex and a demonstration of the segmentation strategy is viewed at far right. Scale bar = 250 μm. (**D**) Sankey plot of all areas of interest segments that passed quality control. (**E**) Principal component analysis plot demonstrating the ability of PCA 1 and 2 to segregate NeuN and pSyn segments, with different regional segments clustering together.

Using this methodology, we collected 103 segments that passed quality control from 11 mice (**Fig. 1D**). Most segments were collected from the ACA and MOs, with MOp having fewer neuronal inclusions, and therefore fewer high-quality segments. Segments were filtered, and segments that did not meet the following criteria were removed: for sequence stitching < 80%, aligned sequence reads < 75%, sequence saturation < 50%, and <5% of genes being detected above the limit of quantification (LOQ). In addition, of the 20175 gene targets contained in the GeoMx Mouse Whole Transcriptome Atlas, 9035 were included in the downstream analyses following QC analysis. Genes were removed based on a global outlier (Grubbs test, p<0.01), local outliers (Grubbs test, p<0.01), and limit of quantification assessment (**Supplementary Fig. 2**). The NeuN and pSyn segments were the major drivers in a principal component analysis (**Fig. 1E**), with cortical layer and region showing less separation.

### α-Synuclein pathology is enriched in layer 5 intratelencephalic and layer 6b neurons

To assess the ability of the captured transcriptome data to differentiate neuron types, we first assessed differentially expressed genes (DEGs) between layers 5 and 6 in the NeuN segment. We identified many DEGs in ACA (**Fig. 2A**), MOs (**Fig. 2B**), and MOp (**Supplementary Fig. 3A**). Many of the DEGs were previously identified to be layer-enriched, including two of the mostly highly DEGs, *Rprm* (enriched in layer 6) and *Etv1* (enriched in layer 5). Enrichment of gene expression was confirmed by assessing *in situ* hybridization staining patterns established by the Allen Institute^23^ (**Fig. 2C**). We next used a published list of genes enriched in specific cell types^24^ to examine cell-type selective DEGs between NeuN and pSyn segments in different brain regions (**Fig. 2D, Supplementary Fig. 3B, 3C**). We found that cell type selective genes were the minority of DEGs, suggesting that the majority of molecular changes are related to the presence of inclusions. However, among those cell type DEGs were some enriched in resilient neurons (*Cadps2*-L5 pyramidal tract (PT), L6 corticothalamic (CT); *Nefm*-L5 PT neurons), while others were enriched in vulnerable neurons (*Cpne4*-L5 intratelencephalic (IT), L6b; *Rorb*-L5 IT neurons).

**Figure 2.**
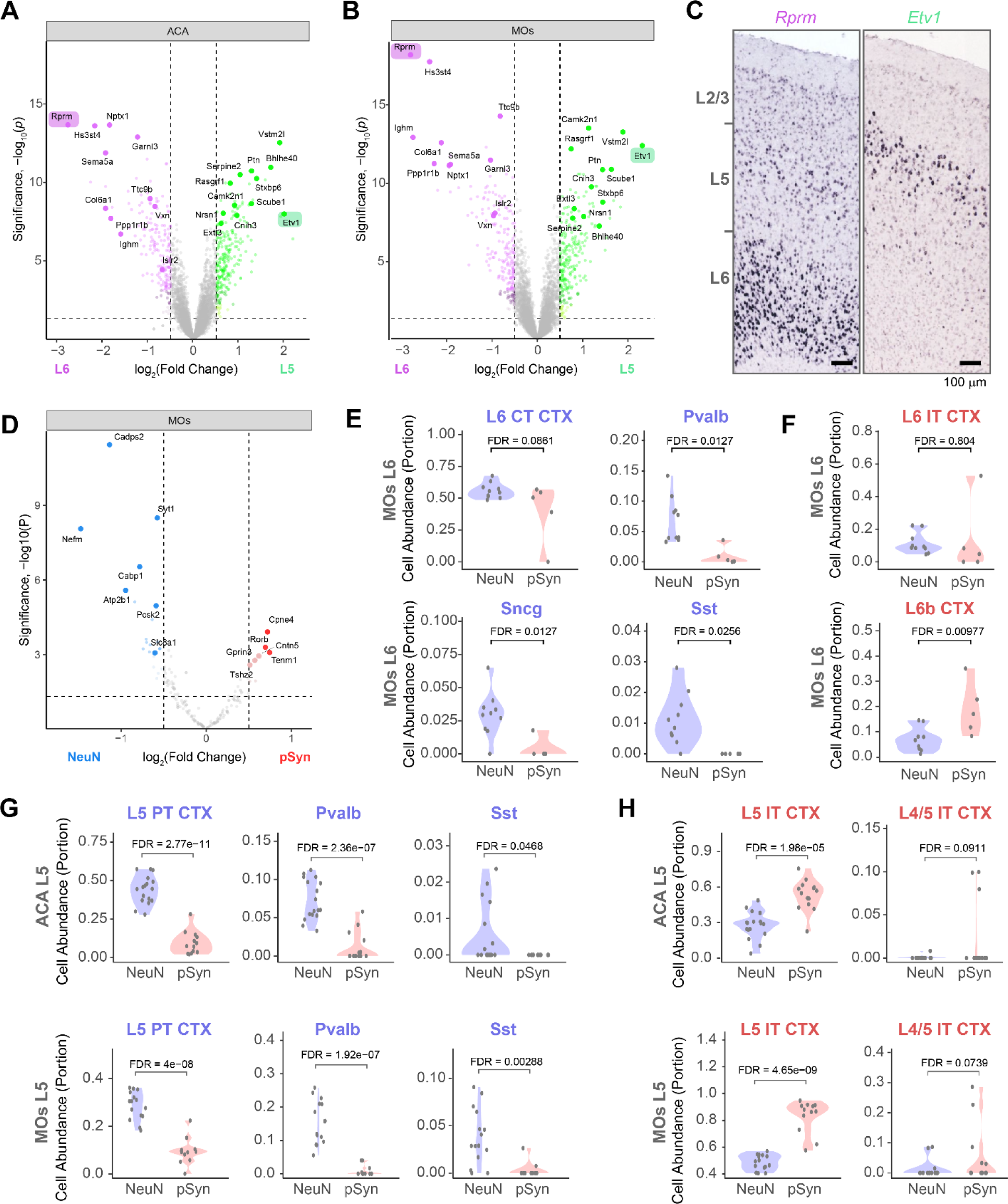
α-Synuclein pathology is enriched in layer 5 intratelencephalic and layer 6b neurons. (**A**) Volcano plot comparing genes differentially expressed between NeuN segments in layer 5 and layer 6 of ACA identifies genes know to be differentially expressed in the different cortical layers. Two such genes are highlighted. (**B**) Volcano plot comparing genes differentially expressed between NeuN segments in layer 5 and layer 6 of MOs. (**C**) *In situ* hybridization images of two genes differentially enriched in layer 6 (*Rprm*) or layer 5 (*Etv1*) of mouse MOs. Images adapted from the Allen Brain Atlas *in situ* data (*Rprm*: https://mouse.brain-map.org/gene/show/43717, *Etv1*: https://mouse.brain-map.org/gene/show/13786). (**D**) Volcano plot comparing genes differentially expressed between NeuN and pSyn segments in the MOs, but only comparing genes known to be differentially expressed by different cell types. (**E**) Cell abundance estimates from cell deconvolution analysis in layer 6 of MOs shows enrichment of Pvalb, Sst, and Sncg cell types with a partial enrichment of L6 CT neurons in resilient neurons. (**F**) Cell abundance estimates from cell deconvolution analysis in layer 6 of MOs show similar proportions of L6 IT neurons and an enrichment of L6b neurons in vulnerable neurons. (**G**) Cell abundance estimates from cell deconvolution analysis in layer 5 of ACA (top) or MOs (bottom) shows enrichment of L5 PT, Pvalb, and Sst neurons in resilient neurons. (**H**) And enrichment of L5 IT and L4/5 IT neurons in the vulnerable neurons. Each dot is an individual segment and FDR-corrected p-values are displayed for statistical comparison.

We next performed cell deconvolution using the SpatialDecon^25^ algorithm to determine cell type proportion using the whole cortex and hippocampus 10X genomics single-cell dataset from Allen Brain Institute as the reference^26^. Clear differences emerged between NeuN and pSyn segments. In layer 6, CT neurons were less abundant in some pSyn segments, and parvalbumin, somatostatin, and *Sncg*+ neuron types were less abundant in all pSyn segments (**Fig. 2E, Supplementary Fig. 4**). L6 IT neurons showed similar abundance in each segment, while L6b neurons were enriched in pSyn segments (**Fig. 2F**). In layer 5, the NeuN segment showed enrichment of layer 5 PT, parvalbumin, and somatostatin neurons (**Fig. 2G**). The pSyn segment showed enrichment of L5 IT neurons and presence of the highly-related L4/5 IT neurons in some segments (**Fig. 2H, 2J**). Thus, we concluded that most inclusions in the cortex of the α-synuclein PFF model at this timepoint are formed in L5 IT, L6 IT, and L6b neurons, while PT and inhibitory neurons are less impacted.

### Excitatory IT neurons are primarily impacted by α-synuclein inclusions

To validate our deconvolution results, we performed cell typing with *in situ* hybridization-immunofluorescence spatially-registered to the Allen Brain Atlas CCFv3. We first stained tissue with a broad excitatory neuron marker (*Slc17a7*), a broad inhibitory neuron marker (*Gad1*) and pSyn (**Fig. 3A**). pSyn inclusions were almost exclusively located in *Slc17a7*+ neurons, while they were absent from *Gad1*+ neurons (**Fig. 3B**, **3C**). To obtain more quantitative data on this localization, we performed cell detection and classification on all neurons in the cortex (**Supplementary Fig. 5**). In these sections, we confirmed that pSyn pathology was primarily in layer 5, with less pathology in layer 6, and minimal pathology in layers 2/3 (**Fig. 3D**). Across layers, almost all pSyn inclusions were in *Slc17a7*+ neurons, while no pSyn inclusions were identified in *Gad1*+ neurons (**Fig. 3E**).

**Figure 3.**
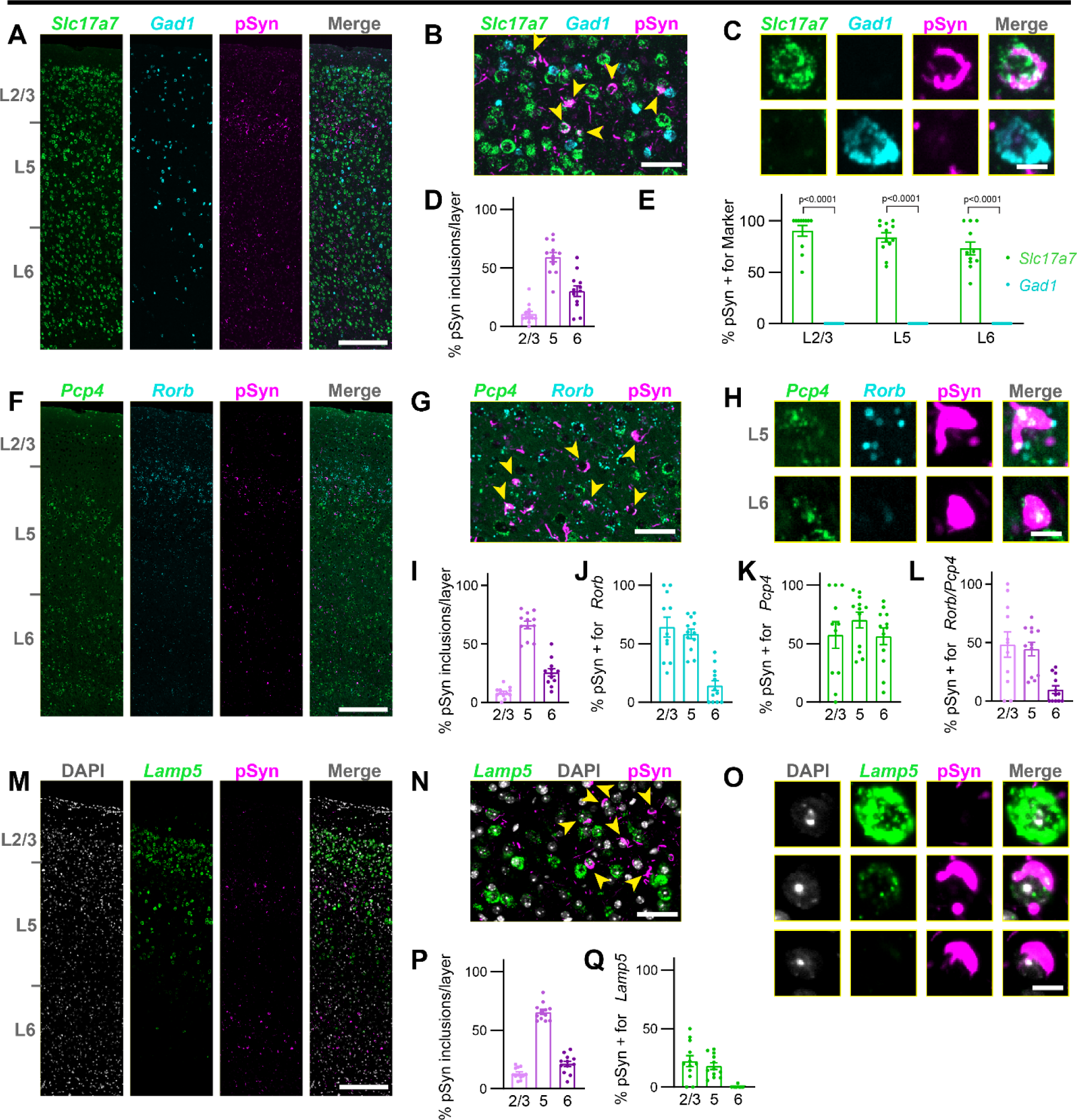
Cell typing, mouse. (**A**) A whole slice through mouse cortex (primarily MOs) stained for *Slc17a7*, *Gad1*, and pSyn. Scale bar = 250 µm. (**B**) A zoomed in view of layer 5 of MOs with yellow arrowheads denoting neurons with pSyn inclusions. Scale bar = 50 µm. (**C**) Examples of single cells positive for *Slc17a7*, *Gad1*, and/or pSyn. Scale bar = 10 µm. (**D**) Quantification of the percentage of pSyn inclusions in each cortical layer as a percentage of the total inclusions shows the highest inclusion burden in L5, followed by L6. (**E**) Quantification of the percentage of cells with pSyn inclusions that were also positive for either *Slc17a7* or *Gad1* in each cortical layer. Plots display the mean +/− SEM with values for individual mice plotted. p-values are displayed from a mixed-effects analysis with Sidak’s multiple comparisons test to compare *Slc17a7* and *Gad1* positive neurons. (**F**) A whole slice through mouse cortex (primarily MOs) stained for *Pcp4*, *Rorb*, and pSyn. Scale bar = 250 µm. (**G**) A zoomed in view of layer 5 of MOs with yellow arrowheads denoting neurons with pSyn inclusions. Scale bar = 50 µm. (**H**) Examples of single cells positive for *Pcp4*, *Rorb*, and/or pSyn. Scale bar = 10 µm. (**I**) Quantification of the percentage of pSyn inclusions in each cortical layer as a percentage of the total inclusions shows the highest inclusion burden in L5, followed by L6. (**J**) Quantification of the percentage of cells with pSyn inclusions that were also positive for *Rorb* in each cortical layer. (**K**) Quantification of the percentage of cells with pSyn inclusions that were also positive for *Pcp4* in each cortical layer. (**L**) Quantification of the percentage of cells with pSyn inclusions that were also positive for both *Rorb* and *Pcp4* in each cortical layer. Plots display the mean +/− SEM with values for individual mice plotted. (**M**) A whole slice through mouse cortex (primarily MOs) stained for DAPI, *Lamp5*, and pSyn. Scale bar = 250 µm. (**N**) A zoomed in view of layer 5 of MOs with yellow arrowheads denoting neurons with pSyn inclusions. Scale bar = 50 µm. (**O**) Examples of single cells positive for *Lamp5* and/or pSyn. Scale bar = 10 µm. (**P**) Quantification of the percentage of pSyn inclusions in each cortical layer as a percentage of the total inclusions shows the highest inclusion burden in L5, followed by L6. (**Q**) Quantification of the percentage of cells with pSyn inclusions that were also positive for *Lamp5* in each cortical layer. Plots display the mean +/− SEM with values for individual mice plotted.

We next sought to identify markers that could more selectively label IT and PT neurons. *Pcp4* is broadly expressed throughout the cortex (**Fig. 3F**) with a preference for excitatory neurons, including both IT and PT neurons in layer 5 and several neuron types in layer 6^26^. Rorb is more selectively expressed, mostly in upper layer 5, bordering layers 2/3 (**Fig. 3F**) and is expressed most highly in IT neurons, with less expression in PT neurons^26^. Most pSyn inclusions in layer 5 were in *Pcp4*+/*Rorb*+ neurons, while neurons in layer 6 lacked *Rorb* staining (**Fig. 3G, 3H**). Quantitatively, pSyn distribution was similar in these sections (**Fig. 3I**) to that previously observed, and layers 2/3 and 5 showed *Rorb* expression in the majority of neurons with pSyn inclusions (**Fig. 3J**). Broad *Pcp4* expression marked most neurons with pSyn inclusions across layers (**Fig. 3K**), so *Rorb*+/*Pcp4*+ quantification mirrors *Rorb*+ alone (**Fig. 3L**).

*Lamp5* is primarily expressed in layer 5 PT neurons and layers 2/3 IT neurons, with lower expression in layer 5 IT neurons (**Fig. 3M**)^26^. Looking particularly in layer 5, pSyn inclusions do not appear in neurons expressing high *Lamp5* (**Fig. 3N**), although a small amount of *Lamp5* can be observed in some inclusion bearing neurons (**Fig. 3O**). Though there are few pSyn inclusions in layers 2/3 (**Fig. 3P**), some of those are in *Lamp5*+ neurons, while very few pSyn inclusions in layer 5 are positive for *Lamp5* (**Fig. 3Q**), and these are in lower-expressing *Lamp5* neurons. Together, these experiments support that pSyn inclusions primarily impact layer 5 IT neurons in the α-synuclein PFF model.

### α-Synuclein inclusion-bearing neurons show gene expression changes associated with cellular dysfunction

We next sought to identify gene expression changes associated with pSyn inclusions. The preponderance of evidence suggests that neurons with pSyn inclusions undergo cellular dysfunction and cell death, but some researchers have argued that inclusions may be protective^2^, sequestering toxic oligomeric species. We conducted DEG analysis in NeuN versus pSyn segments and identified 2298 genes that were upregulated in pSyn segments and 2412 genes that were downregulated in pSyn segments after false discovery rate correction. Hierarchical clustering of segments was able to clearly separate NeuN and pSyn segments by gene expression patterns (**Fig. 4A**). Some of the top genes upregulated in pSyn neurons were related to complement/cytokine (*Itgam*, *Ccl19*), DNA/RNA integrity (*Rpa2, Ercc4, Utp15, Bcl2, Rtkn2*), among others (**Fig. 4B**, **4C**). Top genes downregulated in pSyn neurons were related to the synapse (*Stx1b*, *Cplx1, Dnajc5*, *Nsf*, *Grin2a, Snap25, App, Syt1, Syp*), mitochondria (*Ndufc2, Atp5d, Ndufv2*), lipid metabolism (*Tecr*), endo-lysosome (*Atp6ap2, Psap, Sort1*), ubiquitin-proteasome (*Rpn1, Psma5, Psma7, Herc3*) and cytoskeleton (*Nefm, Brsk1, Kcnc1, Septin7, Kif21a, Stmn2*) among others (**Fig. 4B**, **4C**). Differential expression also appeared highly reproducible across regions (**Supplementary Fig. 6**).

**Figure 4.**
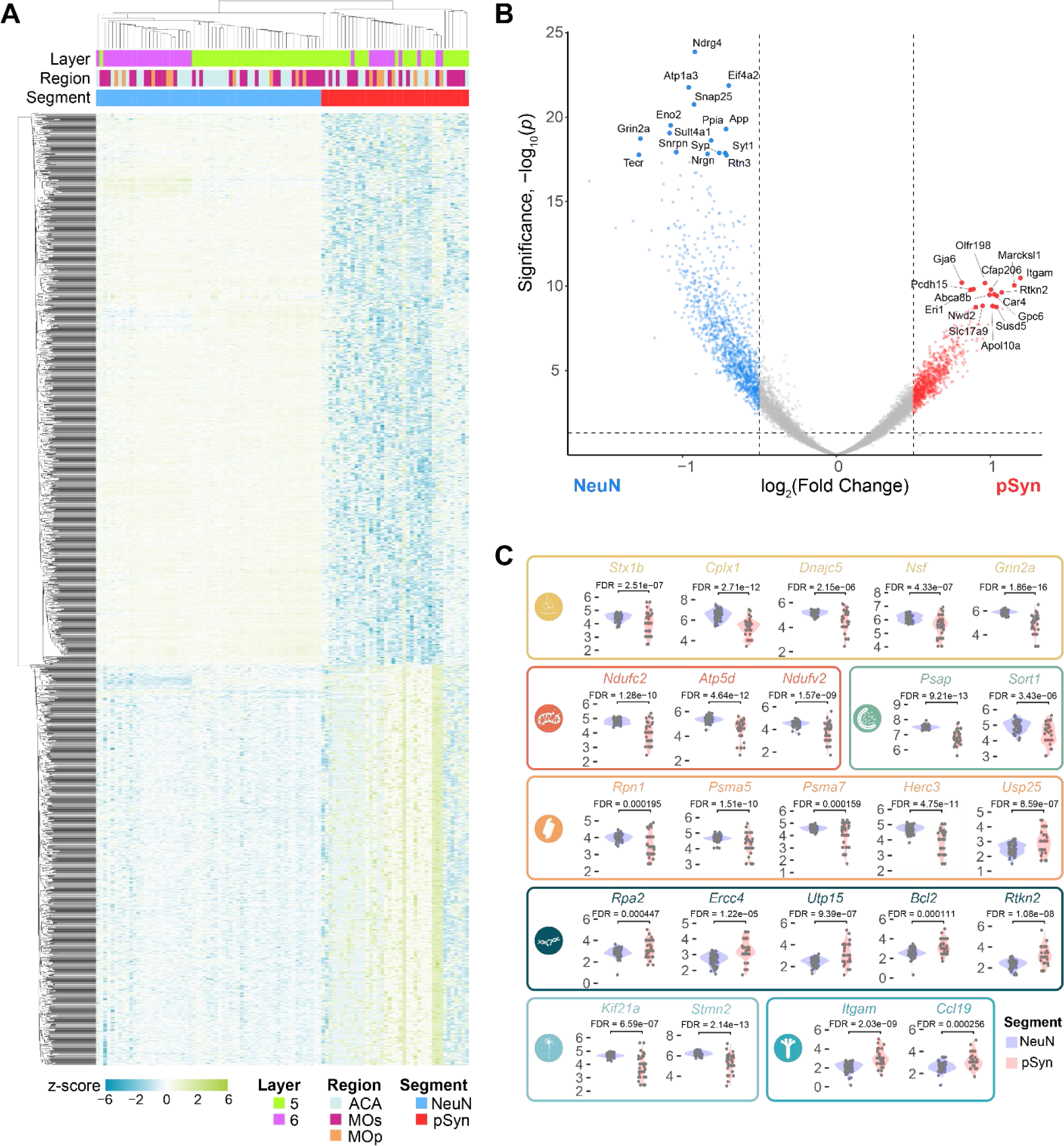
α-Synuclein inclusion-bearing neurons show gene expression changes associated with cellular dysfunction. (**A**) A heatmap displaying the z-score of all genes differentially expressed between pSyn and NeuN segments with FDR-corrected p-value < 0.01. (**B**) Volcano plot comparing genes differentially expressed between NeuN and pSyn segments across regions with the top 30 DEGs labeled. (**C**) Examples of 25 genes that are differentially expressed in NeuN and pSyn segments that fit into one of 9 categories as noted by the color and corresponding symbol. From top-to-bottom, left-to-right, categories are: synapse, mitochondria, lysosome, ubiquitin-proteasome, DNA damage-apoptosis, cytoskeleton, complement-cytokine.

To determine if differentially expressed genes were enriched in particular pathways, we performed gene set enrichment analysis with KEGG Brite pathways^27–29^ comparing pSyn and NeuN segments (**Supplementary Fig. 7**). Pathways downregulated in inclusion-bearing neurons related to the synapse (glutamatergic synapse, dopaminergic synapse, GABAergic synapse, retrograde endocannabinoid signaling, endocytosis), mitochondria (NADH dehydrogenase, NADH:ubiquinone reductase, oxidative phosphorylation), metabolism (citrate cycle), cell signaling (phosphatidylinositol signaling system), RNA (eIF-3, RNA transport), ubiquitin-proteasome (proteasome, family T1 proteasome family, ubiquitin mediated proteolysis), and neurodegenerative disease (Parkinson’s, Alzheimer’s, Huntington’s).

Pathways upregulated in inclusion-bearing neurons related to complement cascade/cytokine (CD molecules, class I cytokine receptor, cytokine-cytokine receptor interaction, complement and coagulation cascades), DNA integrity/apoptosis (apoptosis, small cell lung cancer, Fanconi anemia pathway, helicases), RNA (elongation associated factors), and metabolism (ascorbate and alderate, histidine, fatty acid degradation, hexosyltransferases, cysteine endopeptidases). Together, differential gene expression between NeuN and pSyn neurons revealed that there is broad cellular dysfunction in neurons bearing inclusions with downregulation of pathways essential for neuronal function, and upregulation of stress-related pathways.

### Collection of transcriptome signatures of neurons with or without Lewy pathology with from human cingulate cortex

The α-synuclein PFF model is useful to explore the progressive formation of pathology and neuron death in a homogenous population of mice. LB-like inclusions in mice develop many of the same hallmarks as human LBs^30^, but beyond the inclusions themselves, it is unclear if the cellular phenotype induced resembles PD. We therefore sought to examine neurons with and without LBs in human disease brain (**Fig. 5A**). The cingulate gyrus is one of the first cortical structures to develop Lewy pathology in PD, and one of the most severely affected^31^. It is also functionally analogous to the ACA in mouse that we profiled. We stained the brains of 25 patients diagnosed with PD, PDD or DLB for pSyn, pS202/T205 tau, pS409/410 TDP-43, and amyloid β. All cases were unified Lewy body stage III (brainstem/limbic) or IV (neocortical)^32^ and tau Braak stage IV or below^33^. Of the 25, 13 brains either had minimal Lewy pathology in the cingulate cortex, or also had tau pathology. These cases were not considered for further analysis. The remaining 12 cases showed abundant Lewy pathology, primarily in layer 5, with less pathology in layers 6 and 2/3 (**Fig. 5B**, **5C**).

**Figure 5.**
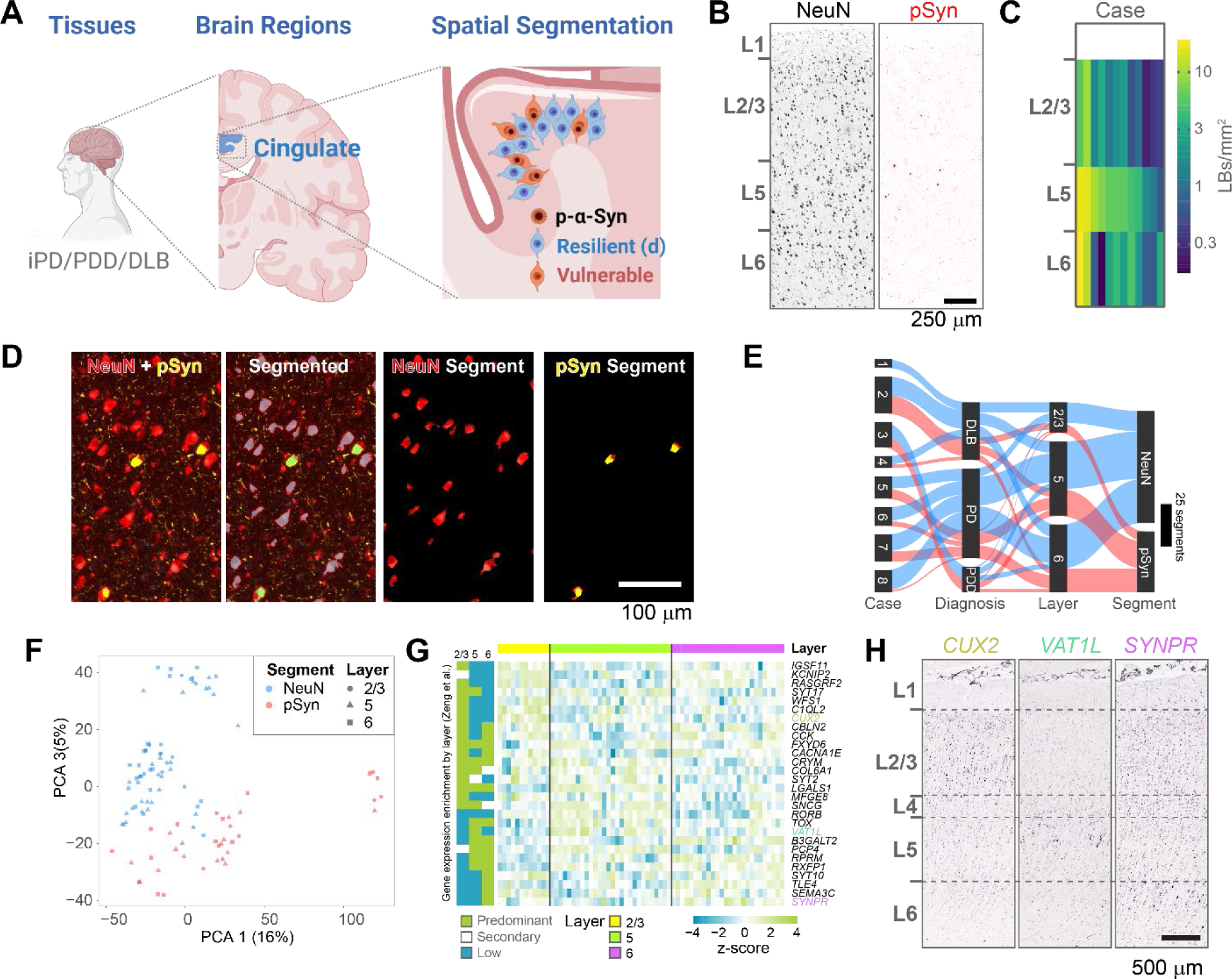
Collection of transcriptome signatures of neurons with or without Lewy pathology with from human cingulate cortex. (**A**) Experimental schematic. Human cingulate cortex tissue was collected from individuals diagnosed with idiopathic PD, PDD, or DLB. (**B**) Example NeuN and pSyn staining from cingulate cortex. Laminar distribution of Lewy pathology can vary between patients with L5 showing the most prominent pathology across cases. Scale bar = 250 μm. (**C**) Quantification of the average number of Lewy bodies per area cingulate cortex used in this study. Layer 5 showed the highest degree of pathology. (**D**) A representative brain section stained for NeuN (red), GFAP (cyan), and pSyn (yellow) with a demonstration of the segmentation strategy. Scale bar = 100 μm. (**E**) Sankey plot of all areas of interest segments that passed quality control. (**F**) Principal component analysis plot demonstrating the ability of PCA 1 and 2 to largely segregate NeuN and pSyn segments, with different cortical layers clustering together. (**G**) Gene expression heatmap for genes significantly differentially expressed by cortical layer in the human GeoMx dataset. Layer specific gene enrichment is noted on the left from Zeng et al.^34^ (**H**) *In situ* hybridization images of three genes differentially enriched in layer 2/3 (*CUX2*), layer 5 (*VAT1L*) or layer 6 (*SYNPR*) of human dorsolateral prefrontal cortex. Images adapted from the Allen Brain Atlas *in situ* data (https://human.brain-map.org/ish/specimen/show/80935564). Scale bar = 500 µm.

Sections of cingulate cortex were stained using the same antibodies as for mouse studies (NeuN, GFAP, pSyn) and imaged on the GeoMx DSP. ROIs were manually drawn within specific cortical layers, and neurons with and without LBs were segmented using NeuN and pSyn fluorescence (**Fig. 5D**). Segments were filtered, and segments that did not meet the following criteria were removed: for sequence stitching < 80%, aligned sequence reads < 75%, sequence saturation < 50%, and <5% of genes being detected above the LOQ. In addition, of the 18815 gene targets contained in the GeoMx Human Whole Transcriptome Atlas, 8601 were included in the downstream analyses following QC analysis. Genes were removed based on a global outlier (Grubbs test, p<0.01), local outliers (Grubbs test, p<0.01), limit of quantification assessment (**Supplementary Fig. 8**). Together, we were able to collect transcript signatures from 8 brains that passed quality control (**Fig. 5E**). Brains were from patients with an average of 78 years old and had an average post-mortem interval of 3.9 hours (**Supplementary Fig. 9**). As expected, data from human brain was substantially more heterogenous than that collected from mice, but the NeuN and pSyn segments were still the main driver of heterogeneity, and a principal component analysis was able to mostly separate the two segments (**Fig. 5F**).

We sought to assess the ability to distinguish cortical layer-selective gene signatures from the NeuN segment. While these signatures are not as well established as they are for mouse cortex, we used the work of Zeng and colleagues to characterize layer-selective gene expression in human cortex^34^ (**Fig. 5G**). We cross-referenced their layer-selective genes with genes in our dataset that showed significant gene expression differences by cortical layer. We identified 28 genes present in both datasets, and the expression patterns were largely replicated (**Fig. 5G**). We evaluated the expression pattern of three of the DEGs by examining in situ hybridization pattens from the dorsolateral prefrontal cortex of Allen Institute cohorts (cingulate cortex was not available). We could see the enrichment of *CUX2* in layers 2/3, *VAT1L* in layer 5, and *SYNPR* in layer 6 (**Fig. 5H**), in agreement with GeoMx expression patterns. These results demonstrate the ability of this method to collect high quality, spatially-resolved transcripts in PD brain.

### LBs are enriched in layer 5 intratelencephalic neurons

In mice, α-synuclein inclusions were primarily in L5 IT neurons and L6b neurons. To assess the distribution of LBs in PD, we carried out analogous staining and cell class quantification experiments (**Supplementary Fig. 10**). We first assessed broad markers of excitatory and inhibitory neuron types, *SLC17A7* and *GAD1*, by *in situ* hybridization-immunofluorescence (**Fig. 6A, 6B, 6C**). LBs in this tissue showed a similar distribution to that seen in α-synuclein PFF-injected mice, with inclusions primarily in layer 5, with lower burden in layers 6 and 2/3 (**Fig. 6D**). Inclusions were also almost exclusively in excitatory *SLC17A7*+ neurons across layers, with few inclusions found in inhibitory *GAD1*+ neurons (**Fig. 6E**).

**Figure 6.**
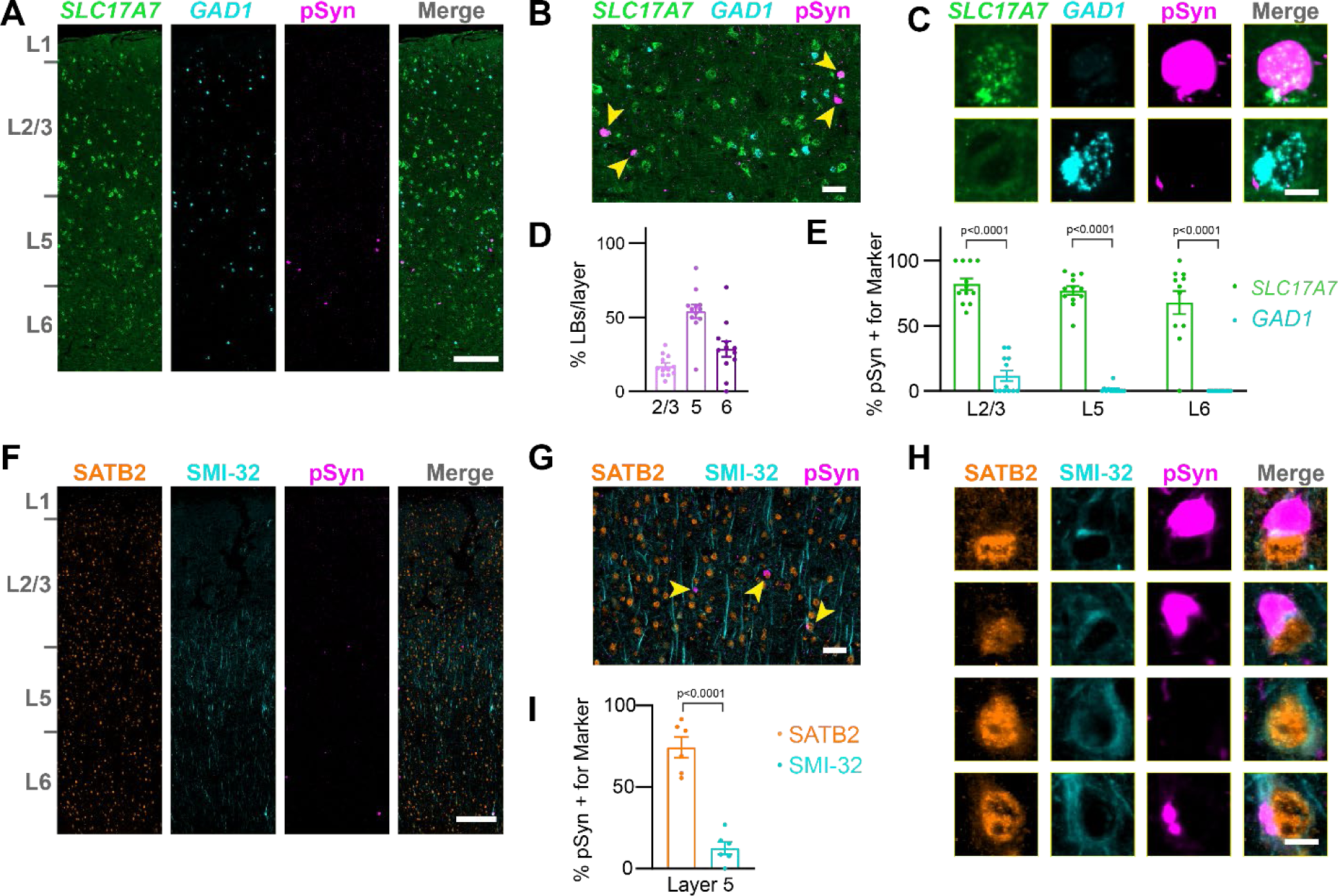
LBs are enriched in layer 5 intratelencephalic neurons. (**A**) A whole slice through human cingulate cortex stained for *SLC17A7*, *GAD1*, and pSyn. Scale bar = 250 µm. (**B**) A zoomed in view of layer 5 with yellow arrowheads denoting neurons with pSyn inclusions. Scale bar = 50 µm. (**C**) Examples of single cells positive for *SLC17A7*, *GAD1*, and/or pSyn. Scale bar = 10 µm. (**D**) Quantification of the percentage of LBs in each cortical layer as a percentage of the total inclusions shows the highest inclusion burden in L5, followed by L6. (**E**) Quantification of the percentage of cells with LBs that were also positive for either *SLC17A7* or *GAD1* in each cortical layer. Plots display the mean +/− SEM with values for individual cases plotted. p-values are displayed from a mixed-effects analysis with Sidak’s multiple comparisons test to compare *SLC17A7* and *GAD1* positive neurons. (**F**) A whole slice through human cingulate cortex stained for SATB2, SMI-32, and pSyn. Scale bar = 250 µm. (**G**) A zoomed in view of layer 5 with yellow arrowheads denoting neurons with pSyn inclusions. Scale bar = 50 µm. (**H**) Examples of single cells positive for SATB2, SMI-32, and/or pSyn. Scale bar = 10 µm. (**I**) Quantification of the percentage of cells with LBs that were also positive for either SATB2 or SMI-32 in each cortical layer. Plots display the mean +/− SEM with values for individual cases plotted. p-value is displayed from a two-tailed t-test to compare SATB2 and SMI-32 positive neurons.

To further assess the balance of LBs in IT versus PT neurons in PD, we stained tissues with antibodies recognizing SATB2, a nuclear protein expressed broadly in excitatory neurons, and neurofilament heavy chain (SMI-32), a protein enriched in L5 PT neurons (**Fig. 6F**). While SATB2 was distributed across cortical layers, SMI-32 showed an enrichment in layer 5. Inclusions were almost all in SATB2+ neurons, while SMI-32 positive neurons only rarely had LBs (**Fig. 6G**, **6H**). Cell classification identifying SATB2+/SMI-32-compared to SATB2+/SMI-32+ neurons showed that around 75% of LBs were only positive for SATB2, while around 12% of neurons were also positive for SMI-32 (**Fig. 6I**). Together, these results support that in PD, layer 5 IT neurons are predominantly impacted by LBs.

### LB-bearing neurons show gene expression changes associated with cellular dysfunction

We next sought to identify gene expression changes associated with LBs. We conducted DEG analysis in NeuN versus pSyn segments and identified 1422 genes that were upregulated in pSyn segments and 620 genes that were downregulated in pSyn segments after false discovery rate correction. Hierarchical clustering of segments was able to clearly separate NeuN and pSyn segments by gene expression patterns (**Fig. 7A**). Some of the top genes upregulated in pSyn neurons were related to complement/cytokine (*C4BPB, IL12RB2*), DNA/RNA integrity (*POLK, TOP3A, AIFM1, GADD45A, NSUN3*), and ion channels (*P2RX6, SCNM1*), among others (**Fig. 7B**, **7C**). Top genes downregulated in pSyn neurons were related to the synapse (*SYNGR1, GRIN1, GRIA2, SYNGAP1, KCNT1*), mitochondria (*NDUFV3, NDUFA10, SDHA*), metabolism (*CKB*), membrane signaling (*CACNA2D3, TMEM59L, TMEM106C*), endo-lysosome (*SORT1, CTSB*), ubiquitin-proteasome (*PSMD8, PSMD1, HERC4, UBE2Z*), and cytoskeleton (*SPTBN2, KIF5A, SEPTIN5, PLEC*) among others (**Fig. 7B**, **7C**).

**Figure 7.**
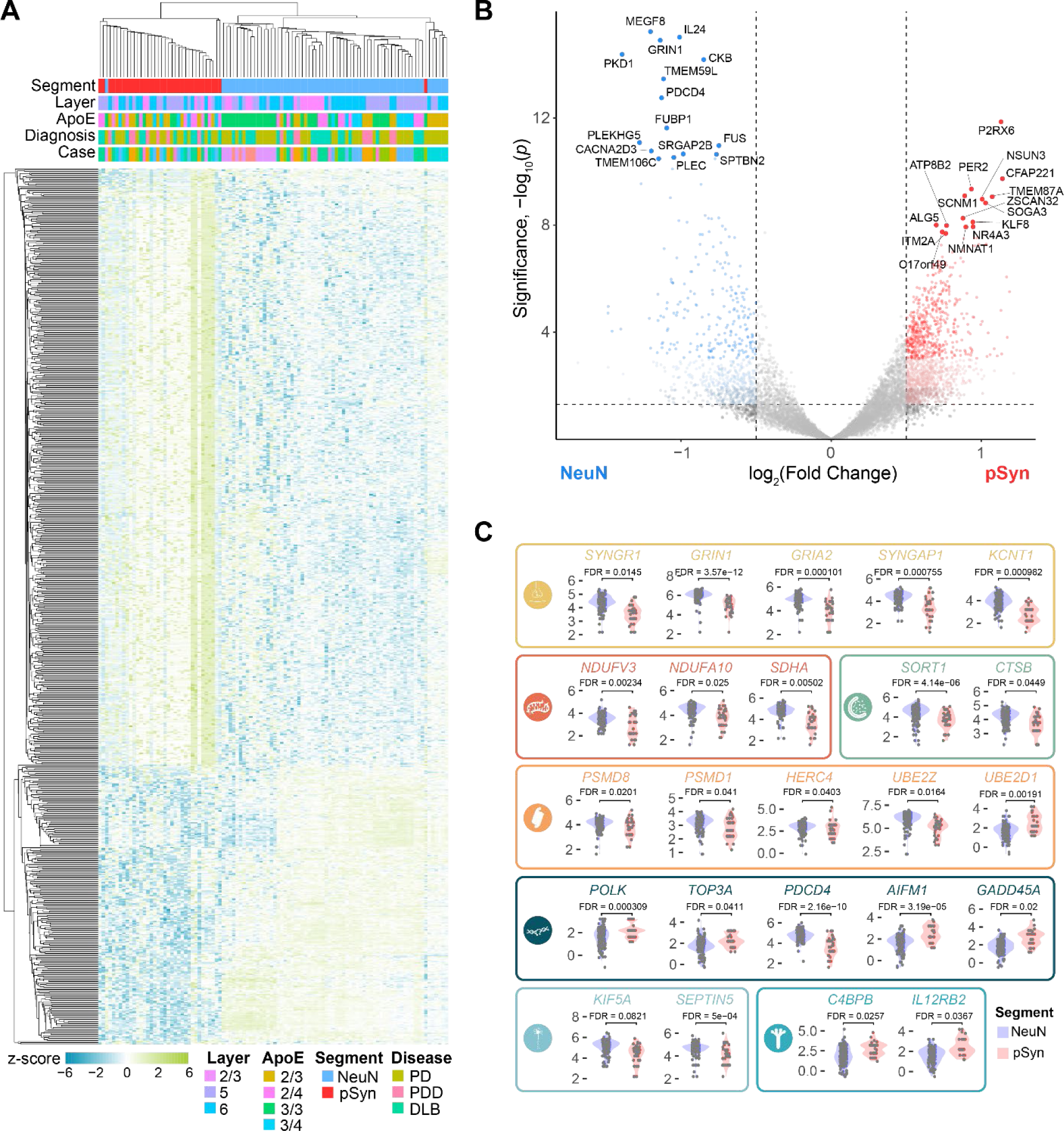
LB-bearing neurons show gene expression changes associated with cellular dysfunction. (**A**) A heatmap displaying the z-score of all genes differentially expressed between pSyn and NeuN segments with FDR-corrected p-value < 0.01. (**B**) (**D**) Volcano plot comparing genes differentially expressed between NeuN and pSyn segments in the across regions with the top 30 DEGs labeled. (**C**) Examples of 25 genes that are differentially expressed in NeuN and pSyn segments that fit into one of 9 categories as noted by the color and corresponding symbol. From top-to-bottom, left-to-right, categories are: synapse, mitochondria, lysosome, ubiquitin-proteasome, DNA damage-apoptosis, cytoskeleton, complement-cytokine.

To determine if differentially expressed genes were enriched in particular pathways, we performed gene set enrichment analysis with KEGG Brite pathways^27–29^ comparing pSyn and NeuN segments (**Supplementary Fig. 11**). Pathways downregulated in LB-bearing neurons related to the synapse (endocytosis, retrograde endocannabinoid signaling), metabolism (sphingolipid signaling pathway), cell signaling (Ca^2+^/calmodulin-dependent kinase, Wnt signaling pathway), RNA (mRNA surveillance pathway, spliceosome, RNA binding proteins, hnRNP proteins, ribosome, small subunit), and neurodegenerative disease (Parkinson’s). Pathways upregulated in LB-bearing neurons related to complement cascade/cytokine (class II cytokine receptor, cytokine-cytokine receptor interaction, complement and coagulation cascades), DNA integrity/apoptosis (base excision repair, mismatch repair), and metabolism (steroid biosynthesis, n-glycan biosynthesis, histidine metabolism, ascorbate and aldarate, tryptophan). Together, differential gene expression between NeuN and pSyn neurons revealed that there is broad cellular dysfunction in neurons bearing inclusions with downregulation of pathways essential for neuronal function, and upregulation of stress-related pathways.

### Parkinson’s disease brain and mice have conserved signatures of α-synuclein inclusions

The α-synuclein PFF model and human tissue each have benefits and weaknesses. The mouse model is highly reproducible on a conserved genetic background, but neurons develop inclusions partially based on injection location. Human brain directly represents PD but individuals have heterogenous genetic backgrounds and experience different environmental insults. To develop a clearer understanding of Lewy pathology related changes, we investigated pathways and gene expression changes that were conserved in mice and humans. 321 and 293 conserved genes were down- and upregulated, respectively, in inclusion-bearing neurons in mice and humans (**Supplementary Fig. 12**). Due to the conservation of gene expression across different neuron types, a mouse model, and human Lewy body diseases, we consider this to be a Lewy-associated molecular dysfunction from aggregates (LAMDA) signature (**Supplementary Fig. 12**).

We identified 29 gene pathways that were significantly enriched in both mouse (**Fig. 8A**) and human (**Fig. 8B**) brain. Pathways downregulated in LAMDA related to the synapse, mitochondria, ubiquitin-proteasome, endo-lysosome, and cytoskeleton, while pathways upregulated in LAMDA related to DNA/RNA integrity and complement/cytokine. Conserved genes were identified that represent each of these pathways (**Fig. 8C, 8D**). In addition, we identified conserved cell type markers (*CUX2, DEPTOR, RORB*), PD-related genes (*PINK1, LRRK2, MAPT*), and cell signaling genes (*PAK6, PDE2A*, **Fig. 8C, 8D**). We further explored the expression of genes associated with monogenic forms of PD^35^. In mice, two genes were elevated in inclusion-bearing neurons (*Lrp10, LRRK2*), while six genes were reduced (*Gigyf2, Pink1, Uchl1, Synj1, Dnajc6, Park7*, **Fig. 8E**). There was a general correspondence in the directionality of these changes in human cingulate, but with more heterogenous results (**Fig. 8F**). Expression of two genes (*PINK1, UCHL1*) was reduced, while expression of three genes (*DNAJC13, PLA2G6, LRRK2*) was increased. Together, these results indicate that there are conserved gene expression changes in mouse and human neurons bearing α-synuclein inclusions.

**Figure 8.**
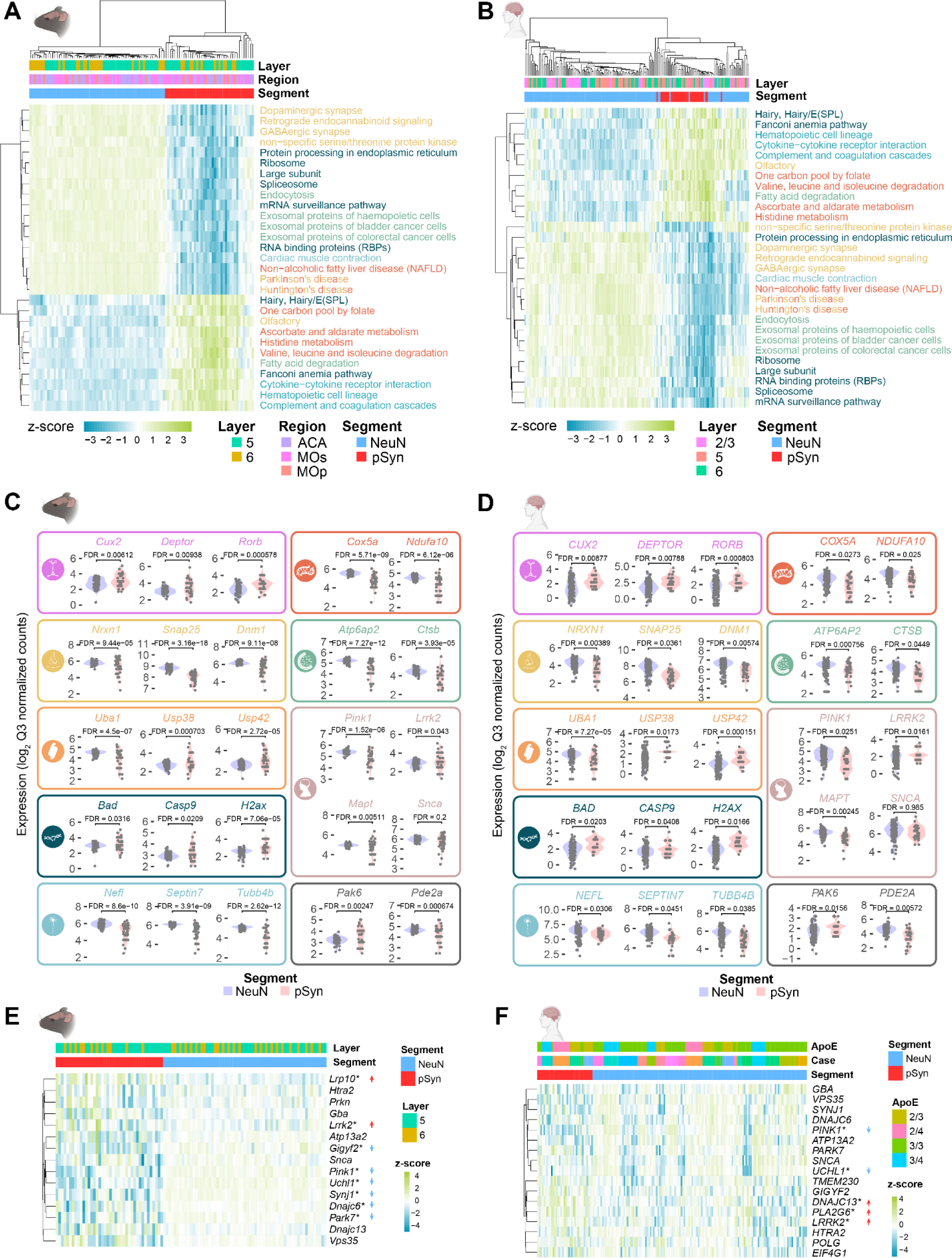
Parkinson’s disease brain and mice have conserved signatures of α-synuclein inclusions. (**A**) Gene set enrichment analysis was performed on NeuN and pSyn segments in from α-synuclein PFF-injected mice. Z-scores of individual segments are plotted for each pathway. Only pathways conserved with human tissue are plotted here. (**B**) Gene set enrichment analysis was performed on NeuN and pSyn segments in from PD/PDD/DLB cingulate cortex. Z-scores of individual segments are plotted for each pathway. Only pathways conserved with mouse tissue are plotted here. (**C**) Representative gene expression from several of the gene sets enriched in mouse NeuN or pSyn segments are plotted. Categories included are (from top to bottom, left to right) cell type, mitochondria, synapse, lysosome, ubiquitin-proteasome, Parkinson’s disease, DNA integrity/apoptosis, cytoskeleton, other. (**D**) Representative gene expression from several of the gene sets enriched in human NeuN or pSyn segments are plotted. Categories included are (from top to bottom, left to right) cell type, mitochondria, synapse, lysosome, ubiquitin-proteasome, Parkinson’s disease, DNA integrity/apoptosis, cytoskeleton, other. (**E**) Expression of genes related to monogenic PD in mice. *FDR-corrected p-value < 0.05. Arrows indicate the direction of significant change. (**F**) Expression of genes related to monogenic PD in human cingulate. *FDR-corrected p-value < 0.05. Arrows indicate the direction of significant change.

## DISCUSSION

The goals of this study were to identify cortical neuron types vulnerable to developing α-synuclein inclusions, and to identify molecular signatures associated with the presence of those inclusions. Soon after the identification of α-synuclein as a main component of LBs^36^, staging studies showed that cortical areas are progressively impacted in PD^16^, and cortical LBs are associated with impaired cognitive performance^4–8^. In the SNc, the ventral tier is most vulnerable to degeneration^11, 12^, especially *SOX-AGTR1*+ neurons^13^. Yet, most knowledge about vulnerability of cortical neurons is based on original staging studies, which found that pathology developed first in deep layers of the cortex and disproportionately affected unmyelinated projection neurons, while myelinated projection neurons (presumably PT) and local circuit neurons, including inhibitory neurons, were protected^16, 37^. These original observations are consistent with our data showing sparing of inhibitory neurons and PT neurons. Interestingly, IT neurons may be selectively vulnerable even in the absence of Lewy pathology. Characterization of the presupplementary motor area in early PD brains with minimal cortical pathology found that cortico-cortical projection neurons were selectively lost in PD, while large SMI-32+ corticospinal neurons were retained^38^.

Excitatory neurons have also been shown to be selectively vulnerable in the α-synuclein PFF model^39^. We expand this work to show that it is selectively L5 IT and L6b neurons. The direct connection of L5 IT neurons to the striatum may be one reason for this vulnerability, but L5 PT neurons also send collateral axons to and through the striatum^40^, giving them the opportunity to be exposed to α-synuclein PFFs as well. Further, the conservation of vulnerability in the α-synuclein PFF model and PD suggest that both connectivity and cell intrinsic vulnerability characteristics may also be conserved.

We also identified conserved molecular changes in aggregate-bearing neurons in both mice and PD that we have designated the LAMDA signature. This first examination of changes within α-synuclein inclusion-bearing neurons demonstrates that these neurons are in a fundamentally altered state where synaptic connections and several cellular functions have been deprioritized and maintenance of DNA integrity or the initiation of apoptosis are prioritized. The LAMDA signature recapitulates the hallmarks of neurodegenerative disease^41^ (pathological protein aggregation, synaptic defects, aberrant proteostasis, cytoskeletal abnormalities, altered energy homeostasis, DNA and RNA defects, neuronal cell death, inflammation), suggesting that even within brains experiencing neurodegeneration, cells bearing aggregates are impacted in a cell-autonomous manner.

Synapses are critical for functional connectivity of neurons, and synapses are impaired early in PD. Dopaminergic terminal loss precedes neuron loss in PD^42, 43^, and this loss of terminals can be used clinically to diagnose PD and monitor progression^44^. It has been proposed that this synaptic vulnerability is related to the burden of maintaining distant, metabolically-demanding presynaptic terminals^45^. Synaptic vulnerability is also seen in glutamatergic neurons. In primary neuron culture, α-synuclein PFF-induced pathology leads to a broad downregulation of synaptic genes and proteins, and a corresponding loss in synapses^46, 47^. In the LAMDA signature, there is broad downregulation of presynaptic genes, especially those involved in the synaptic vesicle cycle (*SNAP25, STXBP1, SYT1, VAMP2, CPLX2, NSF, DNM1*). There is also downregulation of postsynaptic genes including NMDA (*GRIN1*) and AMPA (*GRIA1*) receptors, with the notable exception of some metabotropic glutamate receptors (*GRM7* is elevated in neurons with LBs).

Together with the loss of synapses in PD comes a disruption to axonal transport and the associated cytoskeleton. Previous research has shown that Lewy neurites in the axon may impair axonal cytoskeleton integrity and limit the axonal transport^48^ essential for trophic support and endolysosomal maturation^49^. There is a reduction in cytoskeleton transport proteins early in PD^43^, preceding the loss of TH staining, and α-synuclein pathology has previously been linked to modulation of the actin cytoskeleton^50^. LAMDA includes downregulation of several cytoskeletal genes (*TUBB4B, NEFL, SEPTIN7, CFL1*) and cytoskeleton transport genes (*KIF5A*), consistent with a reduction of synapses and axons in PD.

Mitochondria have also been heavily implicated in PD pathogenesis. α-synuclein fibrils can directly interact with mitochondria, and dysmorphic mitochondria are found within α-synuclein inclusions^46, 51, 52^. This sequestration of damaged mitochondria could be the cause of decreased mitochondrial respiration in primary neurons treated with α-synuclein PFFs^46^. Loss-of-function mutations in PINK1 or Parkin, which control mitophagy, can lead to familial PD^53, 54^ in the absence of Lewy pathology^55^. Interestingly, *PINK1* is expression is decreased in LAMDA, which could be linked to the general downregulation of mitochondrial genes, or it could be a downstream result of LB toxicity. Neurotoxins targeting complex I of the mitochondria, such as MPP+ (1-methyl-4-phenylpyridinium), can induce Parkinsonism by killing dopaminergic neurons^56^ and are therefore used to model PD in rodents^57^. Loss of mitochondrial complex I activity is also sufficient to drive progressive dopaminergic degeneration^58^. Interestingly, the gene deleted in these mice, *Ndufs2*, is downregulated in the LAMDA signature, while a complex I gene whose knockout did not induce degeneration, *Ndufs4*, was not downregulated in LAMDA. It is unclear why oxidative phosphorylation pathways are downregulated in LAMDA. It is possible that metabolic demand on neurons with reduced axons and synapses is reduced, or disruption of mitochondria may precede neuron death, since the mitochondria is intimately linked to the apoptotic process^59^.

Ubiquitination has many functions, but in neurodegeneration it is typically associated with degradation, either via the proteasome, autophagy, or related mitophagy. Protein aggregates, including LBs themselves are highly ubiquitinated^60^. This could be a failed attempt to degrade the misfolded proteins, or a mechanism for clustering them together for further processing^61^. Aggregates are not the only proteins ubiquitinated in neurodegeneration. In particular, ubiquitination of mitochondria is a critical step in mitophagy. Parkin, an E3 ligase, mediates this process after phosphorylation by PINK1. This process is highly controlled by deubiqutinases, including USP30^62^. For that reason, upregulation of PINK1-Parkin or inhibition of USP30 have been pursued as a potential therapeutics for PD^63^.

Interestingly, we find downregulation of *PINK1* and increase in *USP30* and other deubiqutinases (*USP38, USP42*) in LAMDA, suggesting the validity of this therapeutic approach for idiopathic PD. Consistent with the reduced machinery for mitophagy, lysosomal proteases, including *CTSA, CTSB, CTSD* are downregulated. However, it is important to note that the changes seen in LAMDA may be protective for PD as well. An efficient method to minimize circuit disruption and reduce cell-to-cell spread may be to pull back from the network, and if necessary, kill the cell to save the circuit.

Along these same lines, an important question for the field is whether anti-apoptotic treatments would be effective in neurodegenerative diseases or simply modify the inevitable dysfunctional molecular changes that culminate in cell death^64^. We see activation of pathways in LAMDA that have been previously implicated in apoptosis in neurodegenerative disorders^64^. We see an increase in pro-apoptotic factors *BAX* (involved mitochondrial outer membrane permeabilization^59^), *CASP9*, and *BBC3* (p53-upregulated modulator of apoptosis). Due to the role of mitochondria in cell death^59^, we considered a known mitochondria-apoptotic pathway and examine gene expression of that pathway in LAMDA. Mitochondrial outer membrane permeabilization leads to mitochondrial DNA release, which interacts with cGAS (*CGAS*, up in LAMDA) which signals through STING to IRF3 and NF-κB (*Nfkb1*, up in LAMDA) and leads to interferon expression (*Ifnar1* up in mice, *IFNAR2* up in human). We also see induction of complement and cytokine pathways in LAMDA, which is consistent with previous research which has found complement proteins in SNc neurons in PD^65–67^. Thus, gene expression profiles in LAMDA support a potential mitochondria-mediated apoptosis in these neurons. However, there is also an increase in the anti-apoptotic gene *Bcl2* in mice, suggesting that there may be a mixture in inclusion-bearing neurons of staving off apoptosis and promoting it.

The induction of these pathways is consistent with the progressive death of neurons bearing inclusions. While it is difficult to track the trajectory of these neurons in humans, live imaging of transgenic mice injected with α-synuclein PFFs has shown the progressive death of cortical neurons bearing inclusions^68, 69^. Apoptotic pathways can be induced by DNA damage, ER stress, oxidative stress, and cytoskeleton alterations. α-Synuclein inclusions are related to the presence of double-strand DNA breaks and the molecular machinery associated with repair of those breaks (γH2AX) both in mouse and human^70^. This genome instability^71^ may be due to oxidative stress in PD neurons^72^. Several components of the DNA repair pathway are upregulated in LAMDA, including *H2AX*, and genes involved in nucleotide excision repair *GTF2H1, GTF2H3, RPA2*. Manipulation of this pathway has also been found to be protective in *C. elegans* with a-synuclein overexpression^73^.

While no other study has profiled the inclusion status of neurons, recent publications have used RNAseq to profile PD neurons, primarily in the SNc. While some of these studies primarily focused on lost neuron types^13, 14^, more recent studies have also examined gene expression changes in PD neurons^15^. Interestingly, there are some conserved gene signatures between PD SNc neurons, and the cortical LAMDA signature. Synaptic genes (*SNAP25, SYT1, NSF, CACNA1A, GRIK2, GRIA4*), cytoskeletal genes (*NEFL, TUBA1B*), and monogenic PD genes (*SYNJ1, UCHL1*) are downregulated in both datasets. However, not all the SN neurons would be expected to have LBs, and we see interesting differences in these datasets as well. While HSF1 activation is present in the SNc, we see the opposite in LB-bearing neurons (downregulation of *HSPA8, HSPA4L, HSP90AA1, HSPH1*). We also see some conserved increase in metallothioneins (*MT1E, MT1M*), but a contrasting downregulation in others (*MT3*).

There are fewer studies of RNAseq in PD cortex. One study, which profiled excitatory neurons in L5 and L6 of posterior cingulate^18^ identified some changes that were conserved in our dataset, including decreased *OXR1*, which responds to oxidative stress, a decrease in cytoskeletal transport gene *KIF21A*, a decrease in synaptic transmission genes, and decreased proteasomal subunits. However, some gene changes were not replicated, which may reflect changes in neurons without LBs (decrease in *FGF9*). An additional study, which performed bulk and scRNAseq from control and PD cingulate cortex^19^ did not explore specific gene expression changes deeply, but found that the number of DEGs was highest in excitatory neurons, and *PINK1* was downregulated in these neurons, consistent with excitatory neurons bearing LAMDA-related changes.

There is only one other study, to our knowledge, that has examined the transcriptional state of neurons bearing aggregates^74^. This study purified neurons with and without tau neurofibrillary tangles (NFTs) from the prefrontal cortex of Alzheimer’s disease (AD). There appears to be some conserved cellular vulnerability between LBs and NFTs, with L5 IT and L6b neurons identified in both studies. However, additional neurons, including a subset of inhibitory neurons, developed NFTs, while LBs did not appear in those neurons. The DEG profile was completely different in NFT neurons than in LAMDA. Synaptic genes were up in NFT neurons, down in LAMDA. Oxidative phosphorylation and mitochondrial dysfunction were not conserved across NFT neurons, and apoptosis genes were only modestly enriched. Specific genes highlighted as increased in NFT neurons (*HSP90AA1, APP, PRNP*) were all decreased in LAMDA. Additional results from scRNAseq experiments in AD are consistent with the contrasting signatures in AD and PD^75^. Several genes that are increased in AD are decreased in LAMDA (*ST6GALNAC6, BIN1, RAB3A*), although downregulation of metabolic genes is more consistent (*CYCS, NDUFA8* down in AD and LAMDA). These results suggest that while there may be some conservation in factors mediating vulnerability to NFTs and LBs, the cellular response to these two cytoplasmic aggregates is distinct. This distinct cellular response may give insight into why the genetic risk factors for AD and PD are different and also indicate that it may be difficult to find therapies that would be effective in both AD and PD.

There are several limitations to the current study. A primary limitation is the lack of single-cell resolution. We suspect that the heterogeneity in gene signatures, especially among neurons bearing inclusions, is due to a variety of cell states that is lost with the admixture of cells required to obtain sufficient sequence saturation in this study. Future studies may identify a series of cell states in neurons bearing inclusions that can be aligned by pseudotime to reconstruct the trajectory neurons take to dealing with LB inclusions. A second limitation is that we focused on a handful of cortical regions. We suspect that the LAMDA signature may be conserved across cell types in different regions based on similar signatures being identified in SNc neurons in PD^15^, however future studies will be necessary to determine conserved and disparate gene expression changes by cell type.

Leaps have been made recently understanding molecular profiles of cells in neurodegenerative disease. Our study expands on those findings, showing for the first time the molecular changes in neurons with LBs in PD. We further show a conservation of gene expression profile in a mouse model of α-synucleinopathy and PD and have designated this conserved signature LAMDA. We expect LAMDA will provide researchers, including ourselves, the ability to probe our pathways of interest and understand how those are impacted in aggregate-bearing neurons. We also hope it will provide insight about which pathways are most amenable to therapeutic intervention. Finally, we hope that this initial study will be a starting point for future studies of neuronal vulnerability across the brain and through disease progression.

## MATERIALS AND METHODS

### Animals

All housing, breeding, and procedures were performed according to the NIH Guide for the Care and Use of Experimental Animals and approved by the Van Andel Institute Institutional Animal Care and Use Committee (IACUC). C57BL/6J mice were purchased from the Jackson Laboratory (000664; RRID:IMSR_JAX:000664). Both male and female mice were used and were 3-4 months old at the time of injection.

### Recombinant α-synuclein preparation

Purification of recombinant mouse α-synuclein and generation of α-synuclein PFFs was conducted as described elsewhere^76, 77^. The pRK172 plasmid containing the mouse *Snca* sequence was transformed into BL21 (DE3) RIL-competent E. coli (Agilent Technologies Cat#230245). A single colony from this transformation was expanded in Terrific Broth (12 g/L of Bacto-tryptone, 24 g/L of yeast extract 4% (vol/vol) glycerol, 17 mM KH_2_PO_4_ and 72 mM K_2_HPO_4_) with ampicillin. Bacterial pellets from the growth were sonicated and the sample was boiled to precipitate undesired proteins. The supernatant was dialyzed with 10 mM Tris, pH 7.6, 50 mM NaCl, 1 mM EDTA overnight. Protein was filtered with a 0.22 µm filter and concentrated using Amicon Ultra-15 centrifugal filter units (Millipore Sigma Cat#UFC901008). Protein was then loaded onto a Superdex 200 column and 5 mL fractions were collected. Fractions were run on SDS-PAGE and stained with InstaBlue (ApexBio B8226) to select fractions that were highly enriched in α-synuclein. These fractions were combined and dialyzed in 10 mM Tris, pH 7.6, 50 mM NaCl, 1 mM EDTA overnight. Dialyzed fractions were applied to the MonoQ column (GE Health, HiTrap Q HP 645932) and run using a linear gradient from 25 mM NaCl to 1 M NaCl. Collected fractions were run on SDS-PAGE and stained with Coomassie blue. Fractions that were highly enriched in α-synuclein were collected and dialyzed into DPBS. Protein was filtered through a 0.22 µm filter and concentrated to 5 mg/mL (α-synuclein) with Amicon Ultra Centrifugal Filters. Monomer was aliquoted and frozen at −80°C. For preparation of α-synuclein PFFs, α-synuclein monomer was shaken at 1,000 rpm and 37°C for 7 days. Conversion to PFFs was validated by sedimentation at 100,000 x g for 60 minutes and by thioflavin T assay.

### Stereotaxic Injections

All surgery experiments were performed in accordance with protocols approved by the IACUC of Van Andel Institute. α-Synuclein PFFs were vortexed and diluted with DPBS to 2 mg/mL and sonicated in a cooled bath sonicator at 9°C (Diagenode Bioruptor® Pico; 10 cycles; setting medium; 30 seconds on, 30 seconds off). Mice were injected when 3-4 months old. Mice were deeply anesthetized with isoflurane and injected unilaterally into the right forebrain targeting the dorsal striatum (coordinates: +0.2 mm relative to Bregma, +2.0 mm from midline, −2.5 mm beneath the dura) Injections were performed using a 10 µL syringe (Hamilton 7635-01, NV) with a 34-gauge needle (Hamilton 207434, NV) injecting 5 µg α-synuclein PFFs (2.5 µL) at a rate of 0.4 µL/minute. After 3 months, mice were perfused transcardially with PBS and 4% paraformaldehyde (PFA), brains were removed and underwent overnight fixation in 4% PFA. After perfusion and fixation, tissues were processed into paraffin via sequential dehydration and perfusion with paraffin under vacuum (70% ethanol for 1 hour, 80% ethanol for 1 hour, 2 times 95% ethanol for 1 hour, 3 times 100% ethanol for 1 hour, 2 times xylene for 30 minutes, paraffin for 30 minutes at 60°C, paraffin for 45 minutes at 60°C). Brains were then embedded in paraffin blocks, cut into 6 µm sections and mounted on glass slides.

### Human brain tissue

All procedures were done in accordance with local institutional review board guidelines of the Banner Sun Health Research Institute. Written informed consent for autopsy and analysis of tissue sample data was obtained either from patients themselves or their next of kin. Tissues were selected based upon a high burden of Lewy pathology by immunohistochemical staining. Non-pathological cases were balanced by age, sex, and PMI. Human brain tissue was briefly thawed from −80°C on ice before being transferred to ice-cold 10% neutral buffered formalin. Tissue was fixed for 24 hours before being processed into paraffin as noted for mouse tissues. Tissue was cut into 6 µm sections and mounted on glass slides.

### Spatial transcriptomics

Spatial transcriptomics was performed using the nanoString GeoMx® Digital Spatial Profiler. Sections were cut at 6 μm thickness and mounted on plus-charged slides (Epredia Colormark Plus CM-4951WPLUS-001). Slides were baked at 60°C for 1 hour and store at 4°C in a vacuum sealed container containing desiccant for up to two weeks. All subsequent steps were performed using RNase-free conditions and DEPC treated water. Slides were de-paraffinized with 3 sequential 5-minute washes in xylenes, followed by 2 washes in 100% ethanol for 5 minutes, 1 wash in 95% ethanol, and 1 wash in 1x PBS. Target retrieval was performed in target retrieval reagent (10x Invitrogen 00-4956-58 EDTA pH 9.0) diluted to 1x in the BioGenex EZ-Retriever System for 10 minutes at 95°C. Slides were then washed with 1x PBS for 5 minutes. Slides were then incubated in 0.1μg/mL proteinase K (Invitrogen 25530-049) 10 minutes at 37°C and washed in 1x PBS for 5 minutes at room temperature. Slides were post fixed for 5 minutes in 10% neutral buffered formalin followed by two washes in NBF stop buffer (24.5g Tris base and 15g Glycine in 2L DEPC water) for 5 minutes each and one wash in 1x PBS for 5 minutes. Slides were then incubated with hybridization probes (nanoString Cat# 121401103) diluted in Buffer R (provided in the GeoMx RNA Slide Prep FFPE-PCLN kit, catalog # 121300313) in a hybridization oven at 37°C for 16-20 hours.

Following probe incubation, slides were washed with stringent washes (equal parts formamide and 4x SSC buffer) at 37°C twice for 25 minutes each. Then slides were washed twice in 2x SSC buffer. Slides were incubated in 200 μL buffer W (provided in the GeoMx RNA Slide Prep FFPE-PCLN kit, catalog # 121300313) for 30 minutes and incubated in morphology markers (GFAP-488, Thermo 53-9892-82, 1:400; pS129 α-synuclein (81A), Biolegend 825701, 1:1000; NeuN, Millipore ABN78, 1:1000) at 4°C overnight. Slides were washed 4 times in 2x SSC buffer for 3 minutes each wash. Slides were then incubated with secondary antibodies (GαRb 647, Thermo Scientific A21135, 1:1000; GαIgG2a 594, Thermo Scientific A21244, 1:1000) and nuclei marker Syto83 (Thermo Scientific S11364, 1:1000) in Buffer W for 1 hour at room temperature in a humidified chamber. Slides were washed 4 times in 2x SSC buffer for 3 minutes each and placed in the nanoString GeoMx® DSP instrument.

Syto83 immunofluorescence was utilized for autofocus of GeoMx imaging. Immunofluorescence for GFAP was used in identification of morphological markers to aid in fitting to the Allen Brain Institute’s mouse brain atlas. NeuN and pSyn were used for segmentation. Prior to region-of-interest (ROI) generation, the slide image was exported, and individual brains were registered to the Allen Brain Atlas CCFv3 using the mouse brain registration protocol. Images of registered brains were imported onto the DSP instrument and fit exactly to the slide image to enable accurate anatomical selection of ROIs. ROIs were generated using the polygon tool which aligned with cortical layer 5 or layer 6 of the anterior cingulate area (ACAd or ACAv), primary motor cortex (MOp), and secondary motor cortex (MOs). Each ROI was segmented into two areas-of-illumination (AOI): “pSyn” neurons positive for pS129 α-synuclein and NeuN or “NeuN” neurons positive for NeuN and negative for pS129 α-synuclein. Probe identities in each segment were captured via UV illumination and movement to a 96-well plate.

### NGS library prep and sequencing

Illumina Novaseq 6000 was used for sequencing with a read length of 27 for both reads with reverse sequence orientation in the readout group plate information. Plates were dried down and rehydrated in 10 µL nuclease-free water, mixed and incubated at room temperature for 10 minutes. PCR was performed on samples as described in the nanoString GeoMx® DSP Readout User Manual using 2 µL PCR master mix, 4 µL primer from the correct wells, and 4 µL resuspended DSP aspirate. KAPA beads (KAPA Pure beads, Roche Cat# 07983298001) were warmed to room temperature for 30 minutes. Libraries were pooled and KAPA beads were added to each pool at a 1.2X ratio to the final pool volume. Two KAPA bead clean ups were performed and pooled libraries were eluted in 24 µL elution buffer. Negative and positive control pools were eluted into 10 µL elution buffer. Quantity of the pools were assessed using the QuantiFluor® dsDNA System (Promega Corp.). Pools were diluted to 5 ng/µL and quality and size are assessed using Agilent DNA High Sensitivity chip (Agilent Technologies, Inc.) on the Bioanalyzer. Sequencing was performed at 100 reads/µm^2^. Paired end 50 base paired sequencing. Base calling was done by Illumina RTA3 and output was demultiplexed and converted to fastq format with bcl2fastq v1.9.0. Fastq files were then converted to Digital count conversion files (DCC) using GeoMx NGS Pipeline v2.3.3.10.

### *In situ* hybridization-immunofluorescence

In situ hybridization was performed with RNAscope® Multiplex Fluorescent Reagent Kit v2 (ACD, Cat #323270) using recommended conditions and supplied reagents. Paraffin-embedded tissue was freshly sectioned and dried. When it was not going to be used immediately, slides were vacuum-sealed and stored at 4°C. Slide were baked in a dry oven for 1 hour at 60°C and used within one week. Slides were de-paraffinized with 2 sequential 5-minute washes in xylenes, followed by 2 washes in 100% ethanol for 2 minutes. Slides were then dried for 5 minutes at 60°C. For autofluorescence quenching in human brain tissue, slides were incubated in 0.1 M Tris in a humidified tray for 3-5 days under a broad-spectrum LED lamp (Higrow LED, BL-E150A and King LED 3000 watt) until autofluorescence was diminished. Slides were treated with hydrogen peroxide for 10 minutes at room temperature and washed two times with distilled water. Target retrieval was performed in target retrieval reagents in the BioGenex EZ-Retriever System for 15 minutes at 99°C. Slides were then washed with distilled water for 15 seconds and transferred to 100% ethanol for 3 minutes before being dried for 5 minutes at 60°C.

Slides were incubated in protease plus in a humidified tray in a hybridization oven (Boekel Scientific 240200) oven for 30 minutes at 40°C. Slides were washed 2 times with distilled water. RNAscope® probes were added to slides and incubated for 2 hours at 40°C. The following probes were used: *Slc17A7* (Ms:416631, Hu:415611)*, Gad1* (Ms: 400951, Hu:404031)*, Pcp4* (Ms:402311), *Rorb* (Ms:444271), and *Lamp 5*(Ms:451071). Slides were washed twice for 2 minutes with wash buffer and incubated in Amp 1 for 30 minutes at 40°C. The wash and Amp incubation was repeated for Amp 2 and Amp 3, except Amp 3 was only incubated for 15 minutes. Slides were washed twice for 2 minutes with wash buffer and incubated in HRP-C1 for 15 minutes at 40°C. Slides were washed twice for 2 minutes with wash buffer and incubated in Opal 520 (Perkin Elmer FP1487A) or vivid 520 (ACDbio 323271) for 30 minutes at 40°C, washed twice for 2 minutes with wash buffer, incubated in HRP blocker for 15 minutes at 40°C, and washed twice for 2 minutes with wash buffer. The HRP, Opal dye, and HRP blocker steps were repeated for HRP-C2 using Opal 570 (Akoya Biosciences, OP001003) or vivid 570 (ACDbio, 323272). Following the final wash, the slides were further processed for immunofluorescence using the protocol described below, but picking up starting after microwave antigen retrieval.

### Immunohistochemistry/Immunofluorescence

Slides were de-paraffinized with 2 sequential 5-minute washes in xylenes, followed by 1-minute washes in a descending series of ethanols: 100%, 100%, 95%, 80%, 70%. Slides were then incubated in deionized water for one minute prior and transferred to the BioGenex EZ-Retriever System where they were incubated in antigen unmasking solution (Vector Laboratories; Cat# H-3300) and microwaved for 15 minutes at 95°C. Slides were allowed to cool for 20 minutes at room temperature and washed in running tap water for 10 minutes. Slides were incubated in 7.5% hydrogen peroxide in water to quench endogenous peroxidase activity (this step was skipped for immunofluorescence). Slides were washed for 10 minutes in running tap water, 5 minutes in 0.1 M Tris (diluted from 0.5 M Tris made from Tris base and concentrated hydrochloric acid to pH 7.6), then blocked in 0.1 M Tris/2% fetal bovine serum (FBS) for 15 minutes or more. Slides were incubated in primary antibody in 0.1 M Tris/2% FBS in a humidified chamber overnight at 4°C (SMI-32, Biolegend 801702, 1:250, SATB2, Abcam ab92446 1:250, pS129 α-synuclein (81A, Biolegend 825701, 1:1000).

For immunohistochemistry, primary antibody was rinsed off with 0.1 M tris for 5 minutes, then incubated with goat anti-rabbit (Vector BA1000, RRID: AB_2313606) or horse anti-mouse (Vector BA2000, RRID: AB_2313581) biotinylated IgG in 0.1 M tris/2% FBS 1:1000 for 1 hour for immunohistochemistry. Biotinylated antibody was rinsed off with 0.1 M tris for 5 minutes, then incubated with avidin-biotin solution (Vector PK-6100, RRID: AB_2336819) for 1 hour. Slides were then rinsed for 5 minutes with 0.1 M tris, then developed with ImmPACT DAB peroxidase substrate (Vector SK-4105, RRID: AB_2336520) and counterstained briefly with Harris Hematoxylin (Fisher 67-650-01). Slides were washed in running tap water for 5 minutes, dehydrated in ascending ethanol for 1 minute each (70%, 80%, 95%, 100%, 100%), then washed twice in xylenes for 5 minutes and coverslipped in Cytoseal Mounting Media (Fisher, Cat# 23-244-256). Slides were scanned into digital format on an Aperio AT2 microscope using a 20x objective (0.75 NA) into ScanScope virtual slide (.svs) files.

For immunofluorescence, secondary antibodies were incubated on slides in a the dark for 3 hours at room temperature or overnight at 4°C: GαM 488 (Invitrogen A21121, 1:1000), GαRb 546 (Invitrogen A11010, 1:1000), GαIgG2a 647 (Invitrogen A21241, 1:1000). Slides were rinsed twice for 5 minutes in 0.1 M Tris in the dark. Human brain tissue was incubated in Sudan Black (0.3% Sudan Black B (Sigma, Cat# 199664) in 70% ethanol) for 10-120 seconds until autofluorescence was quenched. This step was not performed on RNAscope® slides due to pre-stain autofluorescence quenching. Slides were washed for 5 minutes in 0.1 M Tris and mounted with coverglass in ProLong gold with DAPI (Invitrogen, Cat#P36931). Fluorescent slides were imaged at 20x magnification on a Zeiss AxioScan 7 microscope.

### Cell Detection

Stained slides were scanned on a Zeiss AxioScan 7 at 20x magnification and imported into QuPath for analysis. Individual cells were identified using the Cell Detection feature that allows detection based on DAPI stain intensity. Cell detection parameters such as background radius, sigma, and threshold were adjusted to optimize cell detection across all brain sections.

### Cell Classification

After cell detection, cell classification was completed in QuPath using the object classification feature. QuPath was trained on a subset of annotations to distinguish between different cell types based on signal intensity. To train the classifier, cells were marked as positive or negative for each marker. The training process was repeated for each marker of interest and a composite classifier was created to identify cells positive for multiple markers. Once the classifier was sufficiently accurate, the composite was loaded onto the entire brain image to classify all detected cells. Each cell type was given a unique 6-digit HTML color code.

### Mouse Brain Registration

Images were registered to the Allen Brain Atlas CCFv3 using a modified version of the QUINT workflow^21^. An RGB image of each section was exported from QuPath as a PNG, downsampled by a factor of 12, to use for spatial registration in QuickNII^22^. A segmentation image was created by exporting a color-coded image of classified cells by category on a white background to use as the segmentation input in Nutil. Brain images were uploaded in Filebuilder and saved as an XML file to be compatible with QuickNII. Following the spatial registration of the mouse brain sections to the Allen Mouse Brain Atlas CCFv3 in QuickNII, a JSON file was saved for use in (VisuAlign, RRID:SCR_017978). Brain sections were imported into VisuAlign to fine tune the registrations to match regions of interest. Anchor points were generated in the atlas overlay and moved to the corresponding location on the brain section via non-linear transformations. Markers were placed around the contour of the brain section first with markers refining the inner structures applied second. Final alignments were exported as FLAT and PNG files for use in Nutil^78^.

Nutil was used for the quantification and spatial analysis of the identified cell types in specific regions of the mouse brain. Individual classes were identified for quantification via their HTML color code assigned in QuPath. Nutil generated object counts from each individual classification within each region of the Allen Mouse Brain Atlas using the registration from QuickNII and VisuAlign. Regions of interest, Layers 1, 2/3, 5, 6a, and 6b of the dorsal anterior cingulate area (ACAd), ventral anterior cingulate area (ACAv), primary motor area (MOp), and the secondary motor area (MOs), were extracted from the Nutil output using a custom R code. Furthermore, the dorsal anterior cingulate area (ACAd) and the ventral anterior cingulate area (ACAv) data points were combined (ACA), as well as Layer 6a and Layer 6b for the MOp, MOs, and ACA.

### Statistical analysis

#### Quality Control

In the mouse 3mpi GeoMx experiment, 124 segments were collected in total from 11 mice, and 21 of these segments failed Quality Control (QC) analysis and were removed from the dataset. One segment was removed for sequence stitching < 80% and aligned sequence reads < 75%. Five additional segments were removed for aligned sequence reads < 75% alone, and four segments were removed for sequence saturation < 50%. An additional 11 segments were removed for < 5% of genes being detected above the Limit of Quantification (LOQ). Of the 20175 gene targets contained in the GeoMx Mouse Whole Transcriptome Atlas, 9035 were included in the downstream analyses following QC analysis. 1 gene target was removed as a global outlier (Grubbs test, P<0.01), and 7 were removed as local outliers (Grubbs test, P<0.01). 211 genes were detected below the limit of quantification and were removed. Of the remaining gene targets, 9,031 gene targets were detected above the LOQ in at least 10% of segments, and therefore were selected for further analysis. An additional 4 gene targets were identified as interesting *a priori* and were retained in the study for additional analysis. Detailed QC parameters are described in our mouse analysis pipeline (https://github.com/Goralsth/Spatial-transcriptomics-reveals-molecular-dysfunction-associated-with-Lewy-pathology).

In the human GeoMx experiment 238 segments were collected in total from 8 human PD/PDD/DLB cingulate cortex samples and 11 human healthy control cingulate cortex samples. The Human GeoMx data was assessed on the same parameters as the mouse tissue for Quality Control (QC). In total 63 segments failed QC and were removed from the dataset. 33 samples were removed for low sequence saturation, and 6 segments were removed for low area. An additional 24 segments were removed for < 5% of genes being detected above the Limit of Quantification (LOQ). Of the 18,815 gene targets present in the GeoMx Human Whole Transcriptome Atlas, 8,601 gene targets were included in the downstream analyses following QC. 9 gene targets were removed as local outliers (Grubbs test, P<0.01). 138 genes were detected below the limit of detection and were removed. Of the remaining gene targets, 8,595 gene targets were detected in at least 10% of segments. 6 gene targets were identified as interesting *a priori* and were retained in the study for additional analysis. Detailed QC parameters are described in our human analysis pipeline (https://github.com/Goralsth/Spatial-transcriptomics-reveals-molecular-dysfunction-associated-with-Lewy-pathology).

#### Normalization

To account for systematic variation between AOI’s, we normalized each count by the 75^th^ percentile (Q3) of expression for each AOI. To ensure normalization and Quality Control (QC) procedures were accurate, we compared each AOI’s Q3 value to its limit of quantification. Normalization and QC procedures were determined to be adequate for robust downstream analysis.

#### Differential Expression

Differential expression of gene targets in the GeoMx experiments was determined via a mixed effects linear regression model. The regression model implemented was as follows: y= gene ∼ comparison of interest * variable to control + (1/individual) This model robustly tests for differential gene expression for comparisons of interest (ex. vulnerable vs. resilient neurons, brain region differences, etc.) while controlling for confounding variables and multiple sampling of the same samples.

#### Gene Set Enrichment Analysis

Gene Set Enrichment Analysis was implemented using the *GSVA*^79^ package in R to determine disrupted pathways between vulnerable and resilient neurons. The minimum pathway size was set to 5 genes and maximum pathway size was set to 500 genes. We utilized Kegg Brite pathways to determine which transcriptional pathways are differentially expressed.

#### Cell-type deconvolution

We used nanoString’s cell-type deconvolution algorithm, *SpatialDecon*^25^, to determine the cell-type proportions in our spatial transcriptomics data. We determined cell-type proportions in our data via comparison of our data to single-cell atlas data from the Allen Brain Institute via *SpatialDecon*’s log normal regression. To determine statistical significance of differences in cell type proportions between inclusion bearing (pSyn) and non-inclusion bearing (NeuN) segments, we utilized the following equation: y= cell-type proportion ∼ inclusion status group + (1/individual)

## ACKNOWLEDGEMENTS

We would like to thank the patients and families who participated in this research, without whom this study would not have been possible. We are grateful to the Banner Sun Health Research Institute Brain and Body Donation Program of Sun City, Arizona for the provision of human brains. The Brain and Body Donation Program has been supported by the National Institute of Neurological Disorders and Stroke (U24 NS072026 National Brain and Tissue Resource for Parkinson’s Disease and Related Disorders), the National Institute on Aging (P30 AG19610 and P30AG072980, Arizona Alzheimer’s Disease Center), the Arizona Department of Health Services (contract 211002, Arizona Alzheimer’s Research Center), the Arizona Biomedical Research Commission (contracts 4001, 0011, 05-901 and 1001 to the Arizona Parkinson’s Disease Consortium) and the Michael J. Fox Foundation for Parkinson’s Disease Research.

We thank the Van Andel Institute Genomics Core (RRID:SCR_022913), especially Katelyn Becker, for their assistance with GeoMx spatial transcriptomics. We thank the Van Andel Institute Bioinformatics and Biostatistics Core, especially Zachary Madaj, for their assistance with statistical analysis. We thank the Van Andel Institute Pathology and Biorepository Core (RRID:SCR_022912), especially Lisa Turner, for their assistance with GeoMx staining. We thank the Van Andel Institute Optical Imaging Core (RRID:SCR_021968), especially Corinne Esquibel, for their assistance with slide imaging. This research was supported in part by the Van Andel Institute Vivarium (Grand Rapids, MI).

This research was funded by Aligning Science Across Parkinson’s ASAP-020616 to M.X.H. and NIH grant R01-AG077573 to M.X.H. Several images were created with BioRender.com.

## AUTHOR CONTRIBUTIONS

T.G. designed the experiments, performed experiments, analyzed results, and wrote the manuscript. L.M., L.B., L.B, K.K. performed experiments and analyzed results. D.D. developed analysis tools and analyzed results. L.T., K.B, and M.A. designed and performed experiments. D.N. analyzed results. M.X.H. conceived and designed the experiments, performed experiments, analyzed results, and wrote the manuscript. All authors have reviewed and approved the manuscript.

## MATERIALS & CORRESPONDENCE

Correspondence and material requests should be addressed to M.X.H.

## DATA AVAILABILITY

The data that support the findings of this study are available from data repositories.

## CODE AVAILABILITY

Primary data and code used to generate the analyze gene expression and cell detection is available on GitHub: https://github.com/Goralsth/Spatial-transcriptomics-reveals-molecular-dysfunction-associated-with-Lewy-pathology and https://github.com/DaniellaDeWeerd/NutilToUsable.

---

**Supplementary Fig. 1.**
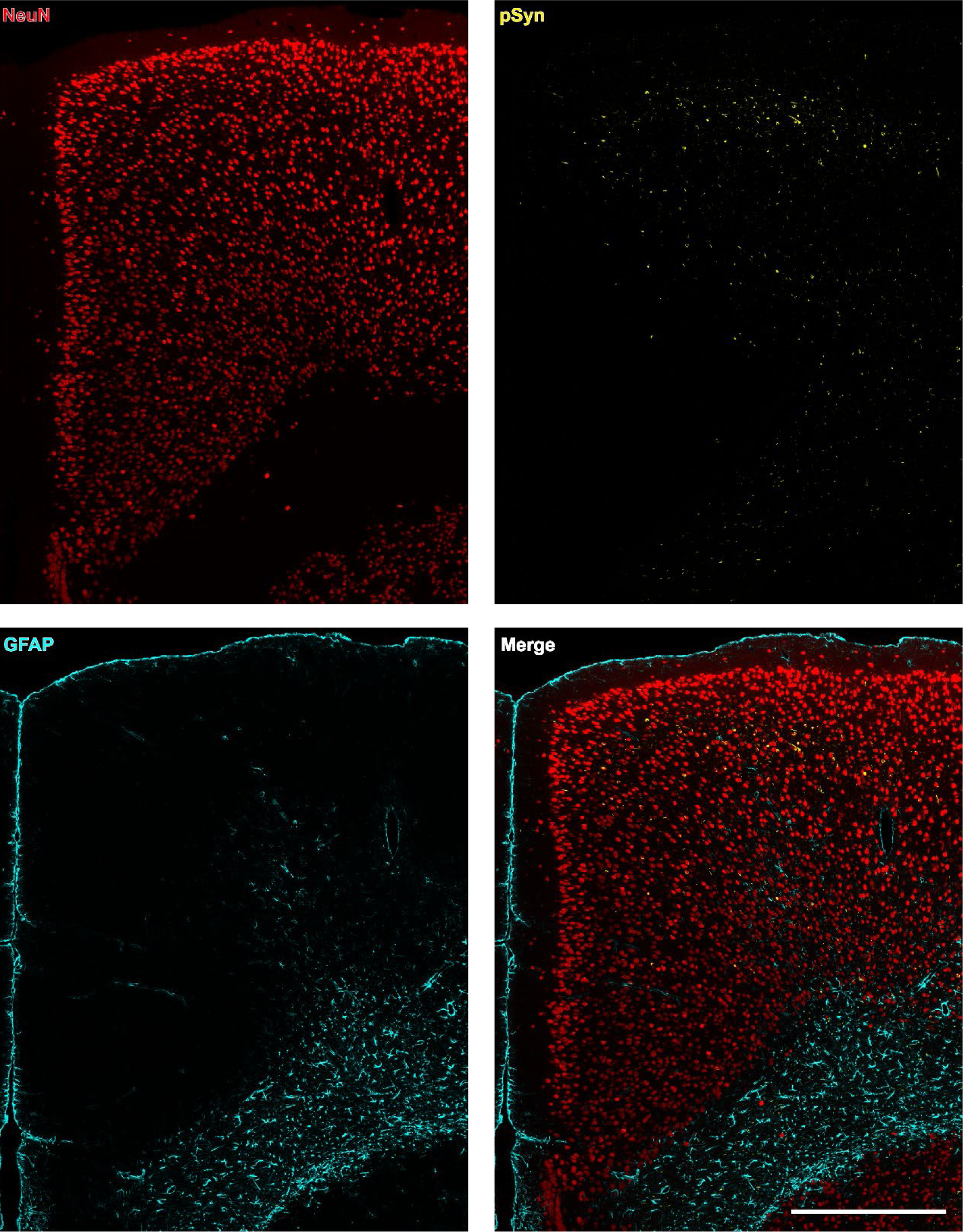
Representative cortex from α-synuclein PFF-injected mouse. Three different proteins were stained for segmentation on the GeoMx DSP instrument. NeuN stains neuron cell bodies. pSyn stains α-synuclein pathology. GFAP stains for astrocytes. pSyn pathology is primarily observed in upper layer 5 and in layer 6 of cortex. Scale bar = 0.5 mm.

**Supplementary Fig. 2.**
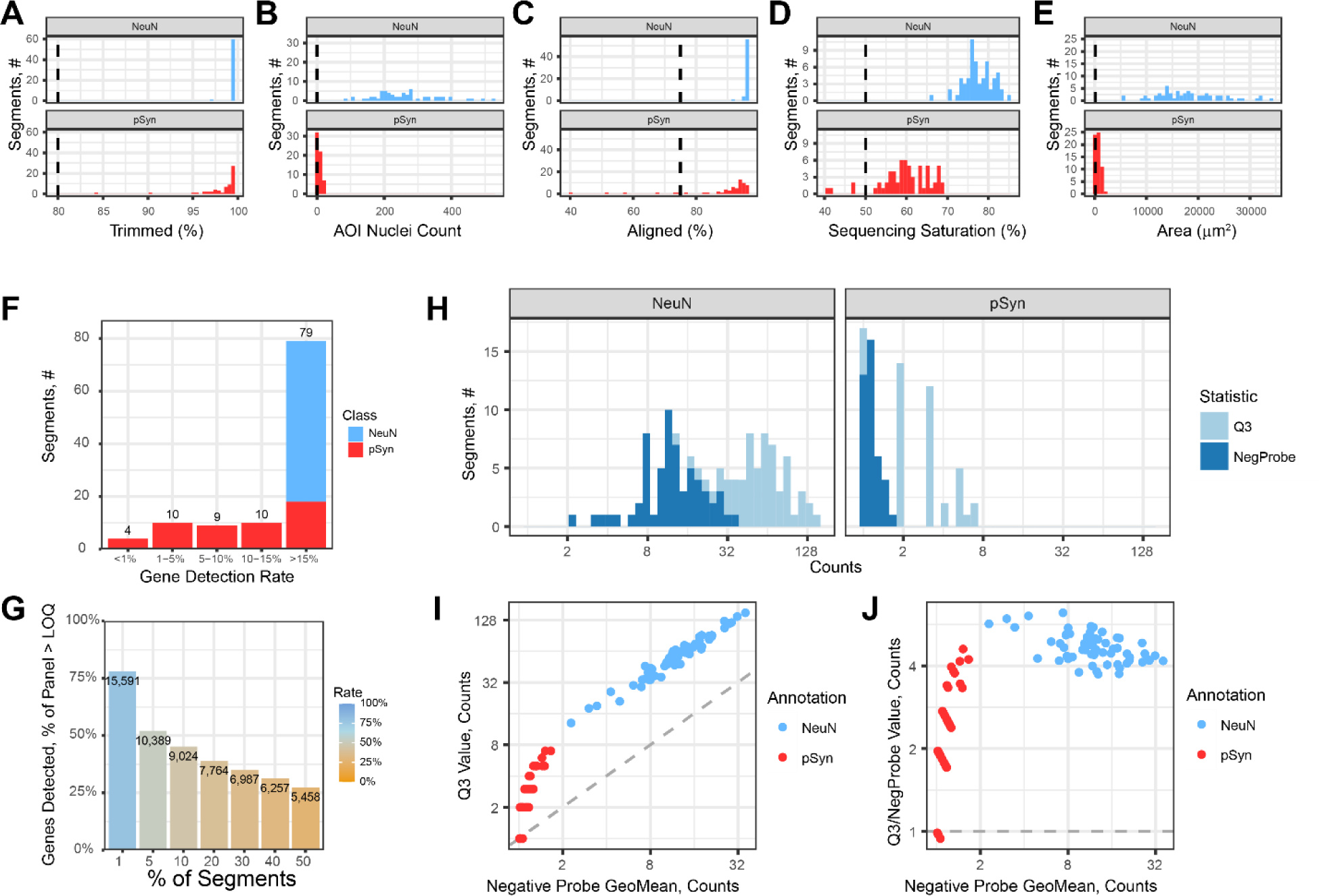
Quality control of mouse GeoMx data. **(A-E)** Assessment of segments for quality based on spatial information and standard transcriptomics quality control metrics. **(F)** Number of segments with the given percentage of genes detected above the limit of quantification (LOQ). **(H)** Assessment of negative probe count values and Q3 values for each segment. **(G)** Assessment of the number of genes detected above LOQ in the given percentage of segments. **(I)** Comparison of the segments Q3 count value to the geometric mean of the negative probes. **(J)** Comparison of the segments Q3 values divided by their negative probes geometric mean to the negative probe geometric mean.

**Supplementary Fig. 3.**
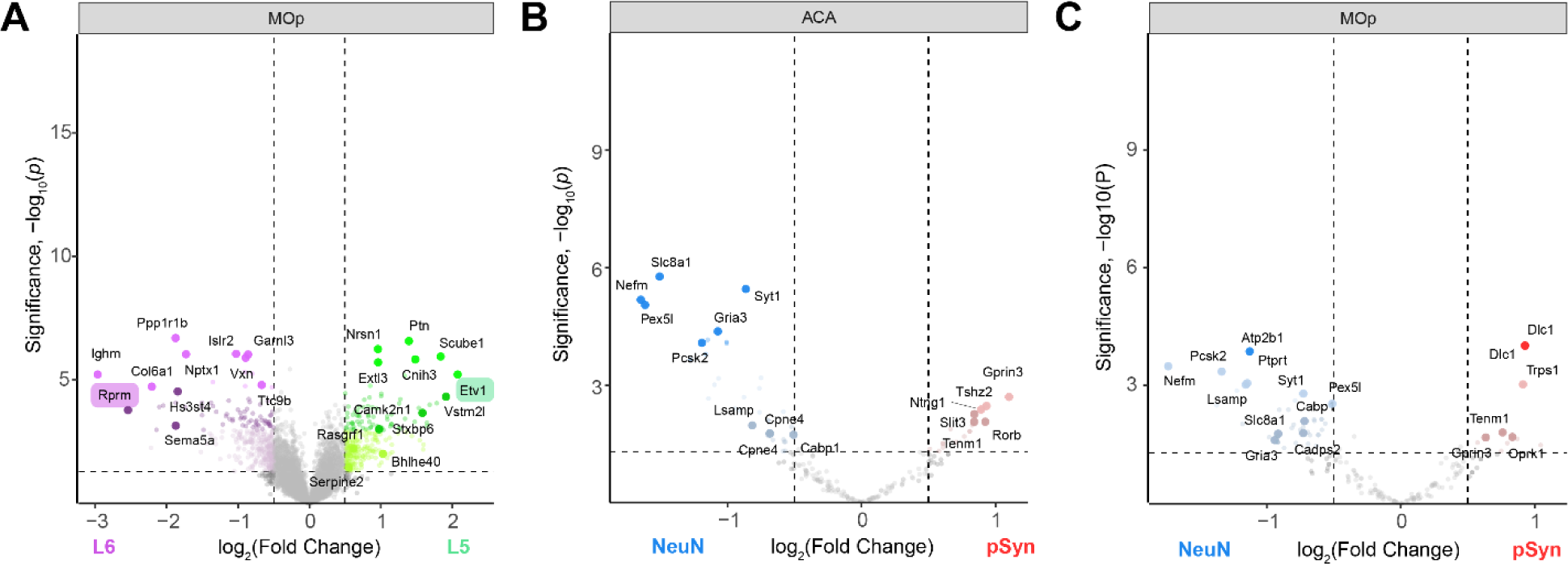
Differential gene expression by layer and cell types. (**A**) Volcano plot comparing genes differentially expressed between NeuN segments in layer 5 and layer 6 of MOp identifies genes know to be differentially expressed in the different cortical layers. Two such genes are highlighted. (**B**) Volcano plot comparing genes differentially expressed between NeuN and pSyn segments in the ACA, but only comparing genes known to be differentially expressed by different cell types. (**C**) Volcano plot comparing genes differentially expressed between NeuN and pSyn segments in the MOp, but only comparing genes known to be differentially expressed by different cell types.

**Supplementary Fig. 4.**
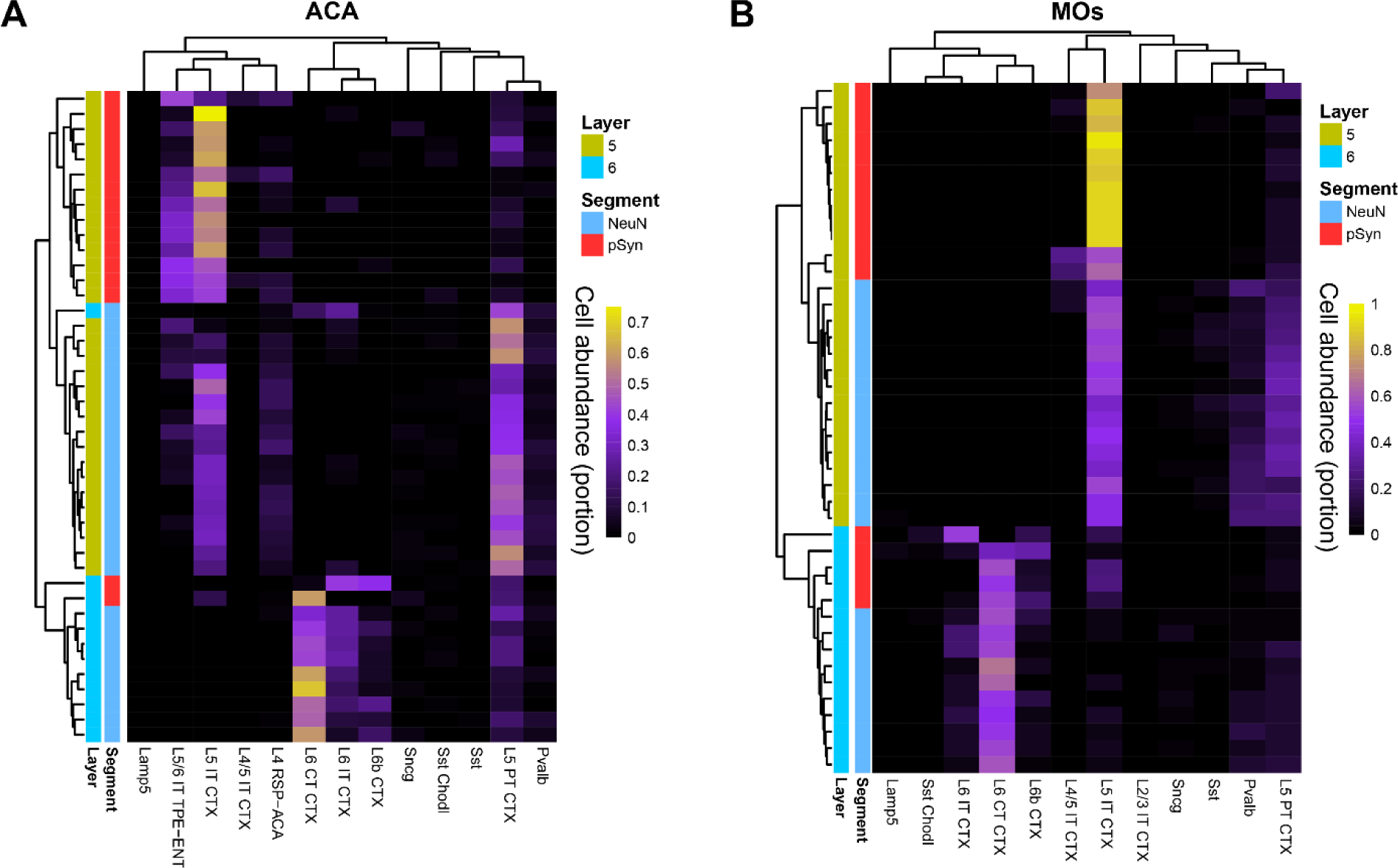
Cell deconvolution plots. (**A**) The relative abundance of cell types was calculated via cell deconvolution for each segment in the ACA region. Layer 5 and layer 6 segments largely cluster separately, with NeuN and pSyn segments clustering within their respective layers. Layer 5 pSyn segments are largely L5 IT or L5/6 IT TPE-ENT cell types, while layer 5 NeuN segments show less abundance of L5 IT and a higher proportion of L5 PT neurons. Layer 6 pSyn segments are a mix of L6 CT and L6b neurons, while layer 6 NeuN segments are mostly L6 CT and L6 IT neurons. Similar results are seen for the MOs region (**B**).

**Supplementary Fig. 5.**
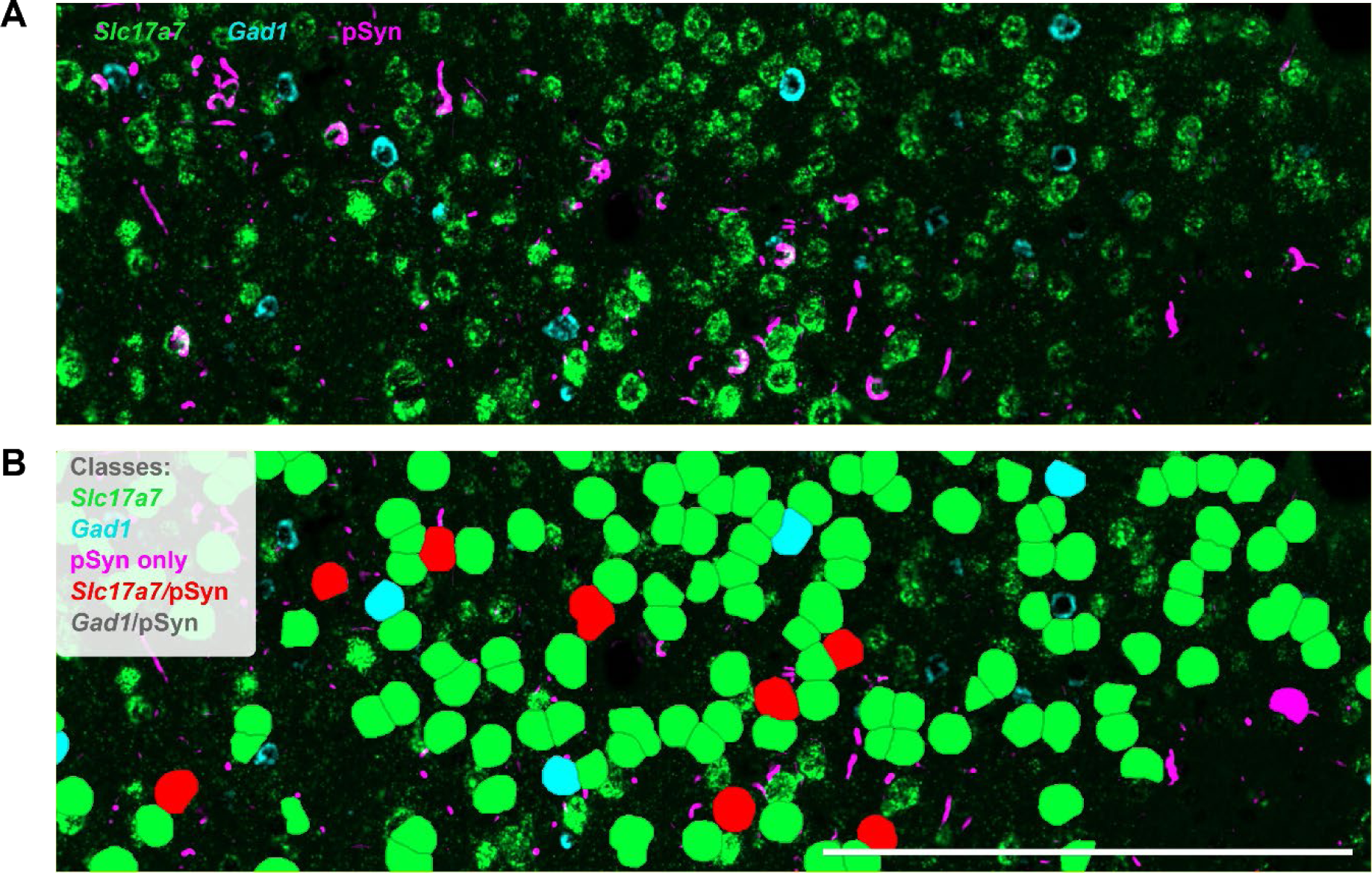
Cell classification in mouse cortex. (**A**) Mouse cortex stained for RNAs *Slc17a7* and *Gad1* and protein pSyn. (**B**) The same image as in panel **A**, except cells were identified based on the presence of nuclear DAPI signal and classified based on the presence or absence of *Slc17a7*, *Gad1*, or pSyn in the cytoplasm. Cells with more than one marker were classified as having multiple markers. The overlay indicates the cell class. Scale bar = 250 µm.

**Supplementary Fig. 6.**
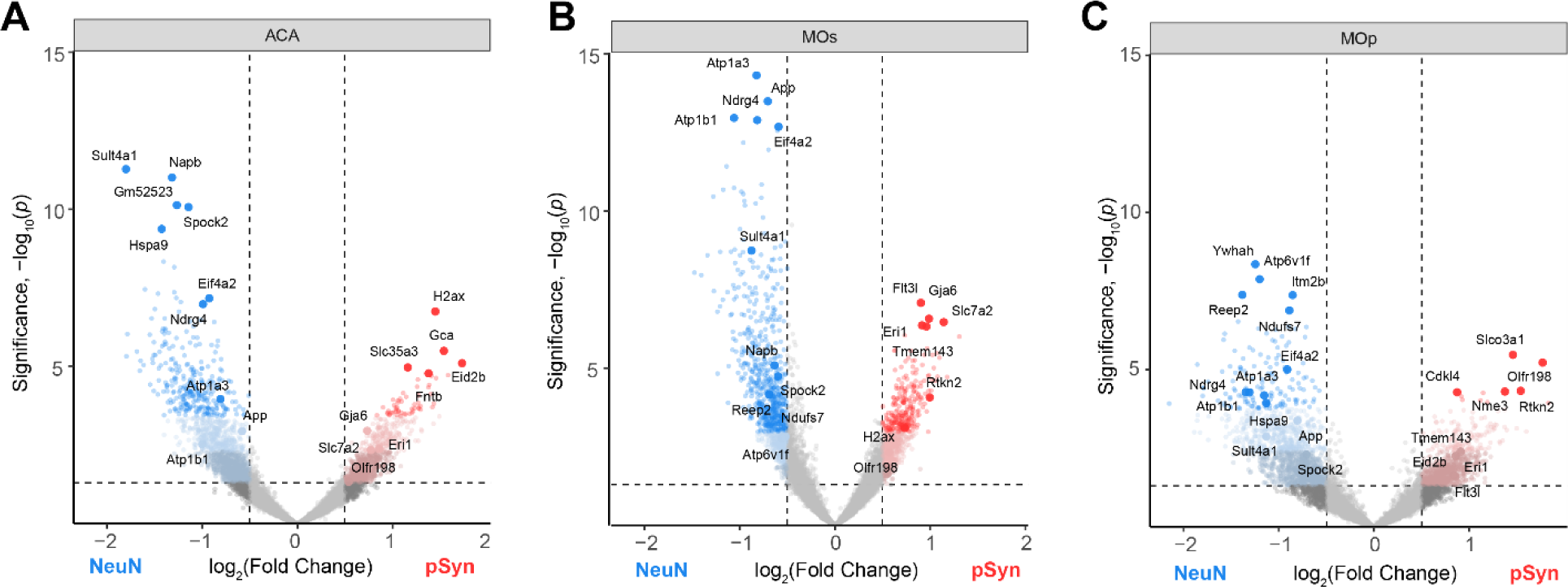
α-Synuclein inclusion-bearing neurons show conserved gene expression changes associated with cellular dysfunction in three cortical regions Volcano plot comparing genes differentially expressed between NeuN and pSyn segments in the ACA (A), MOs (B), or MOp (C) regions with the top 30 DEGs labeled.

**Supplementary Fig. 7.**
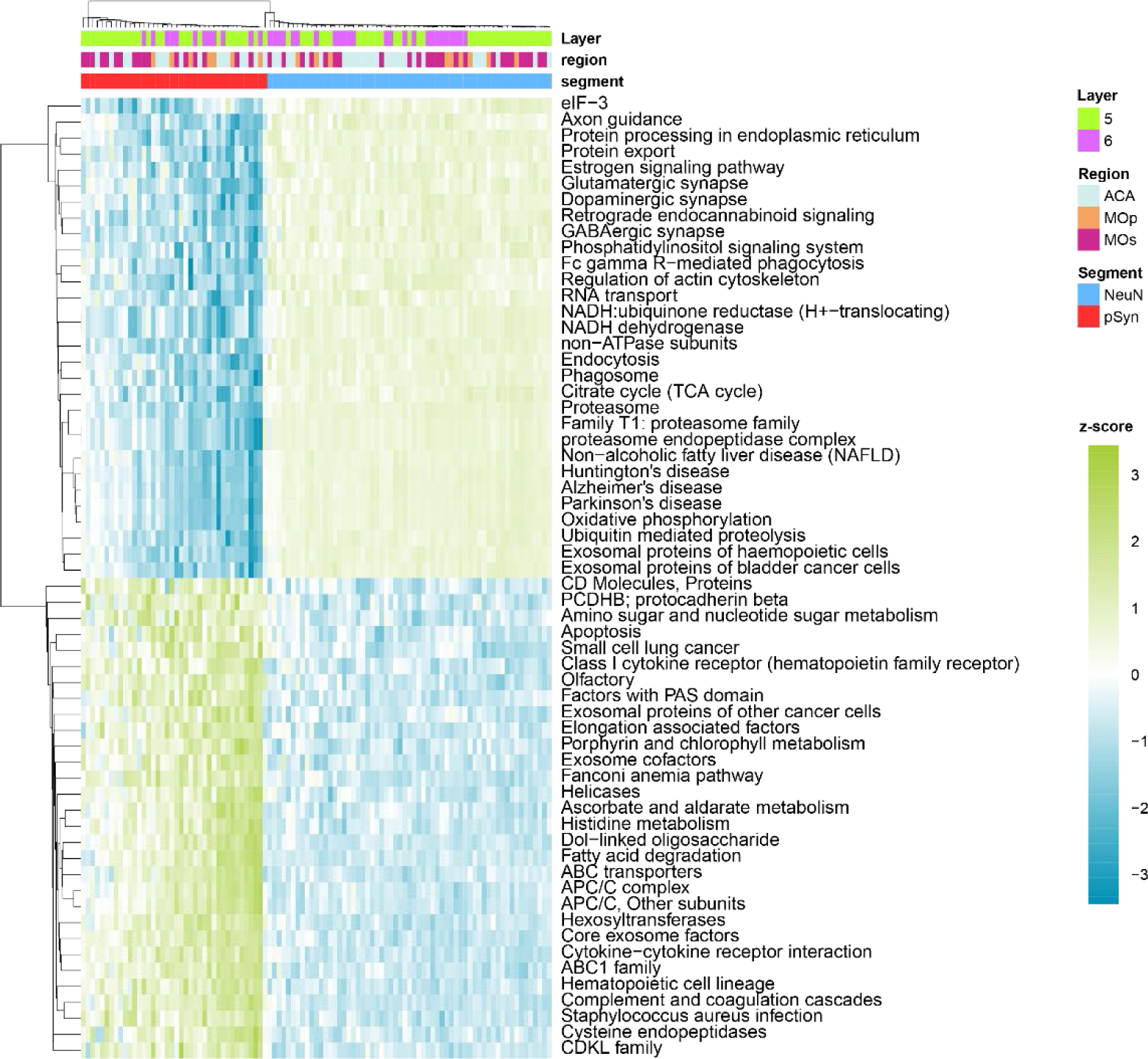
Mouse gene set enrichment analysis. Gene set enrichment analysis was performed on NeuN and pSyn segments in from α-synuclein PFF-injected mice. Z-scores of individual segments are plotted for each pathway. The top 60 pathways enriched in either pSyn or NeuN segments are plotted.

**Supplementary Fig. 8.**
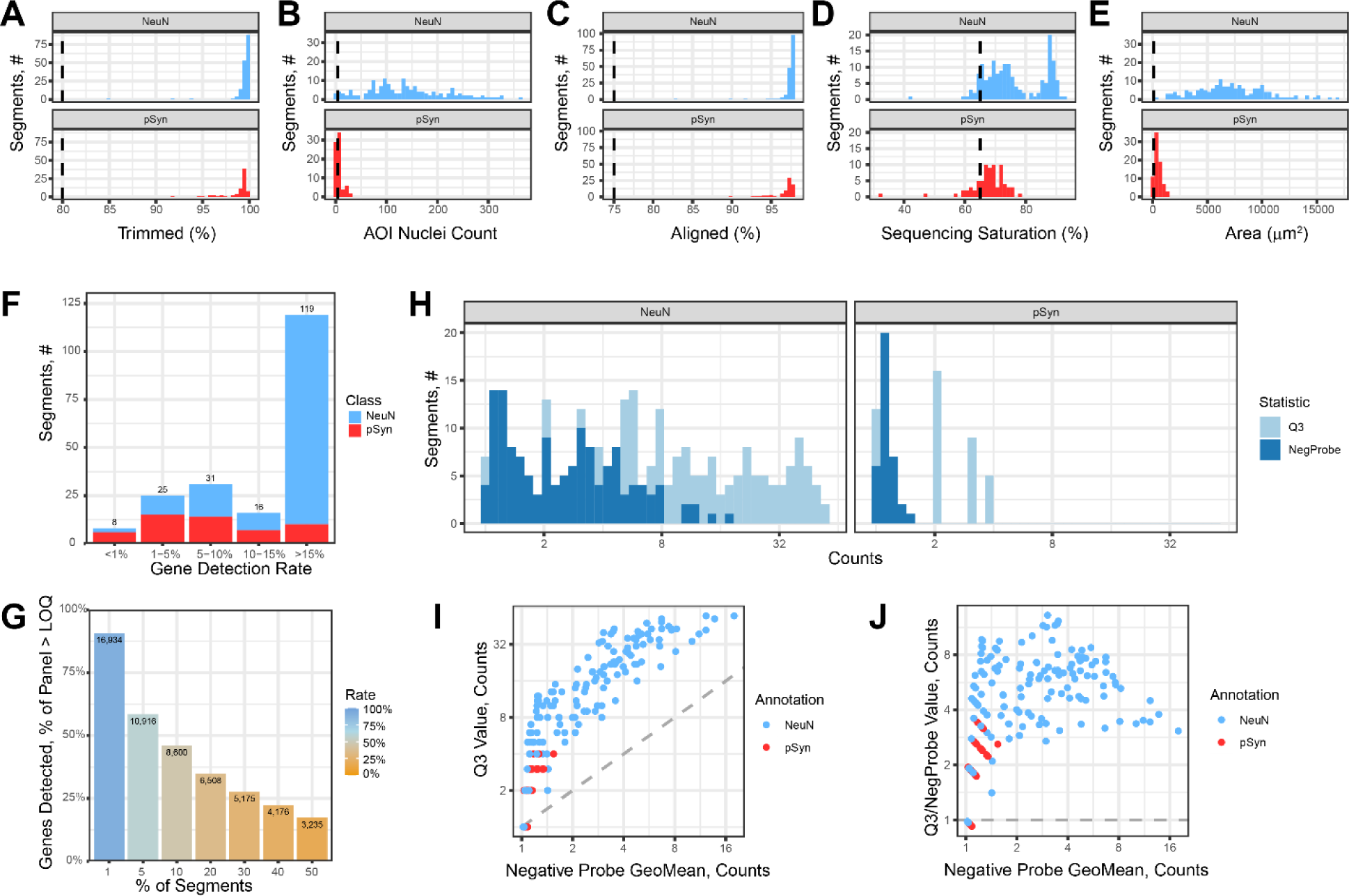
Quality control of human GeoMx data. **(A-E)** Assessment of segments for quality based on spatial information and standard transcriptomics quality control metrics. **(F)** Number of segments with the given percentage of genes detected above the limit of quantification (LOQ). **(H)** Assessment of negative probe count values and Q3 values for each segment. **(G)** Assessment of the number of genes detected above LOQ in the given percentage of segments. **(I)** Comparison of the segments Q3 count value to the geometric mean of the negative probes. **(J)** Comparison of the segments Q3 values divided by their negative probes geometric mean to the negative probe geometric mean.

**Supplementary Fig. 9.**
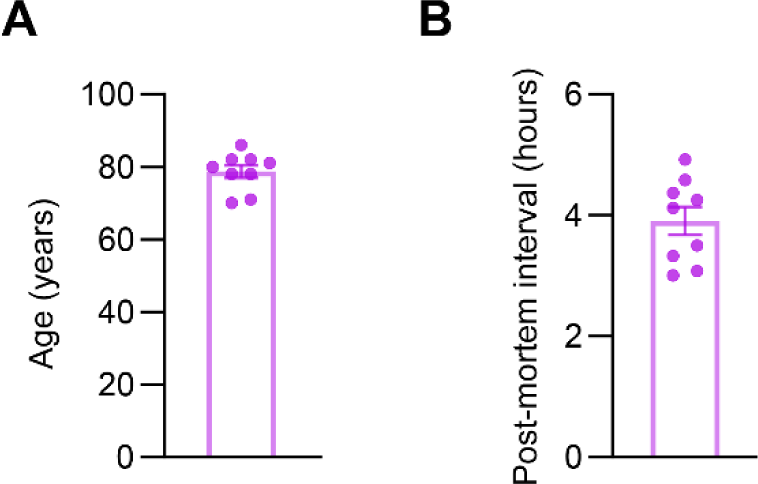
Human brain tissue information. (**A**) Age and (**B**) post-mortem interval is shown for PD, PDD, and DLB cases used for GeoMx analysis.

**Supplementary Fig. 10.**
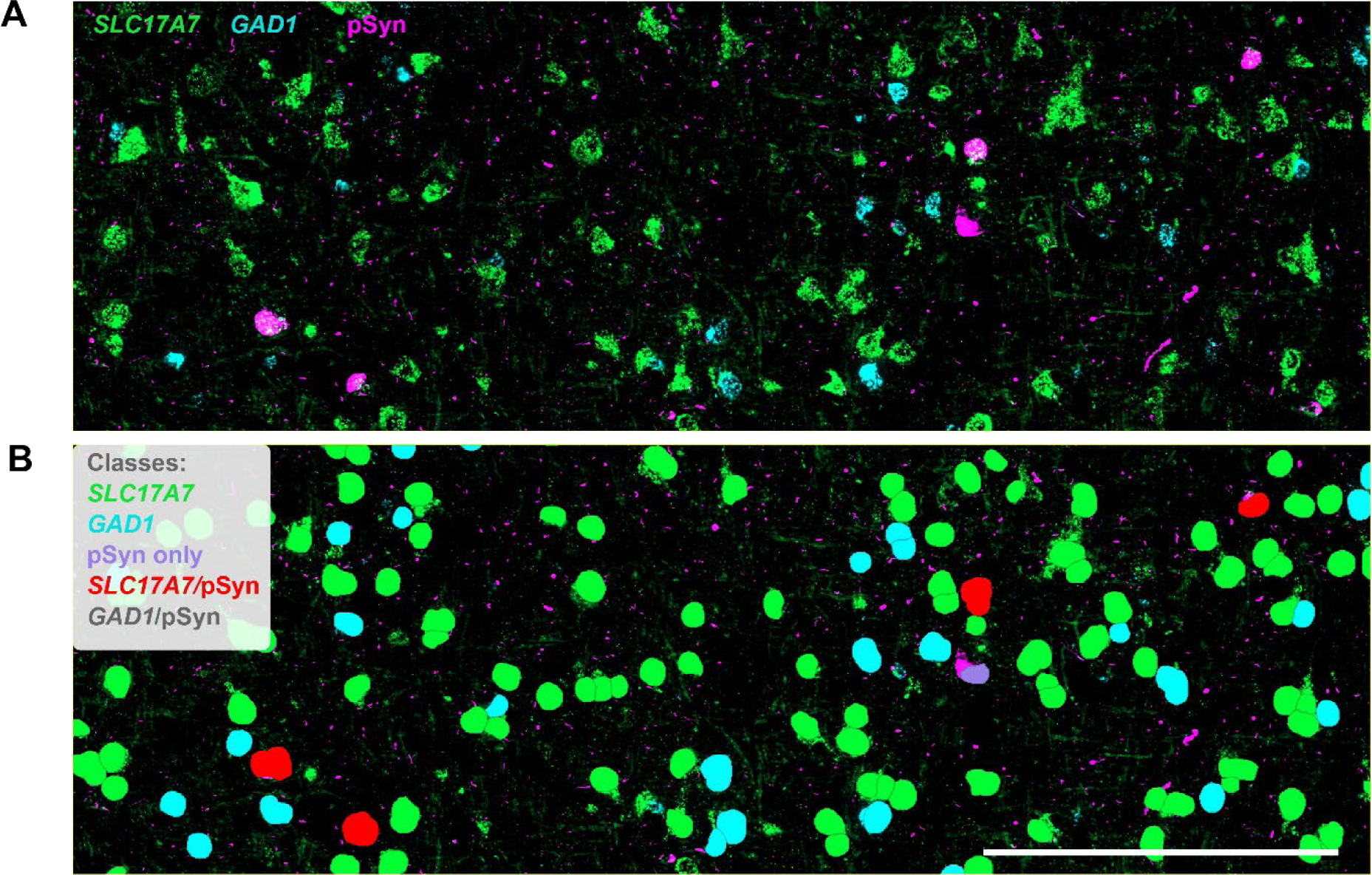
Cell classification in human cortex. (**A**) Human cingulate cortex stained for RNAs *SLC17A7* and *GAD1* and protein pSyn. (**B**) The same image as in panel **A**, except cells were identified based on the presence of nuclear DAPI signal and classified based on the presence or absence of *SLC17A7*, *GAD1*, or pSyn in the cytoplasm. Cells with more than one marker were classified as having multiple markers. The overlay indicates the cell class. Scale bar = 250 µm.

**Supplementary Fig. 11.**
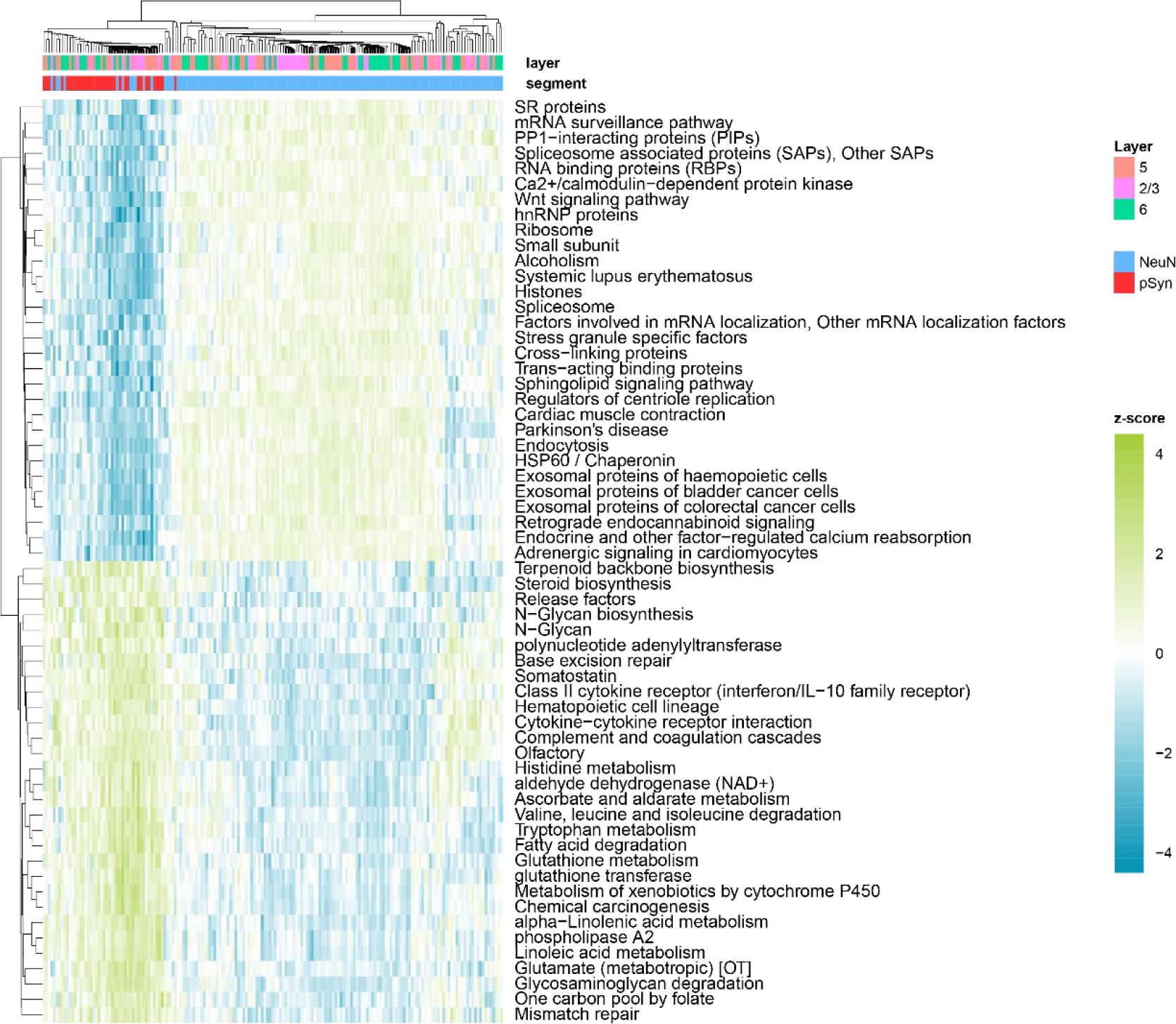
Human gene set enrichment analysis. Gene set enrichment analysis was performed on NeuN and pSyn segments in from α-synuclein PFF-injected mice. Z-scores of individual segments are plotted for each pathway. The top 60 pathways enriched in either pSyn or NeuN segments are plotted.

**Supplementary Fig. 12.**
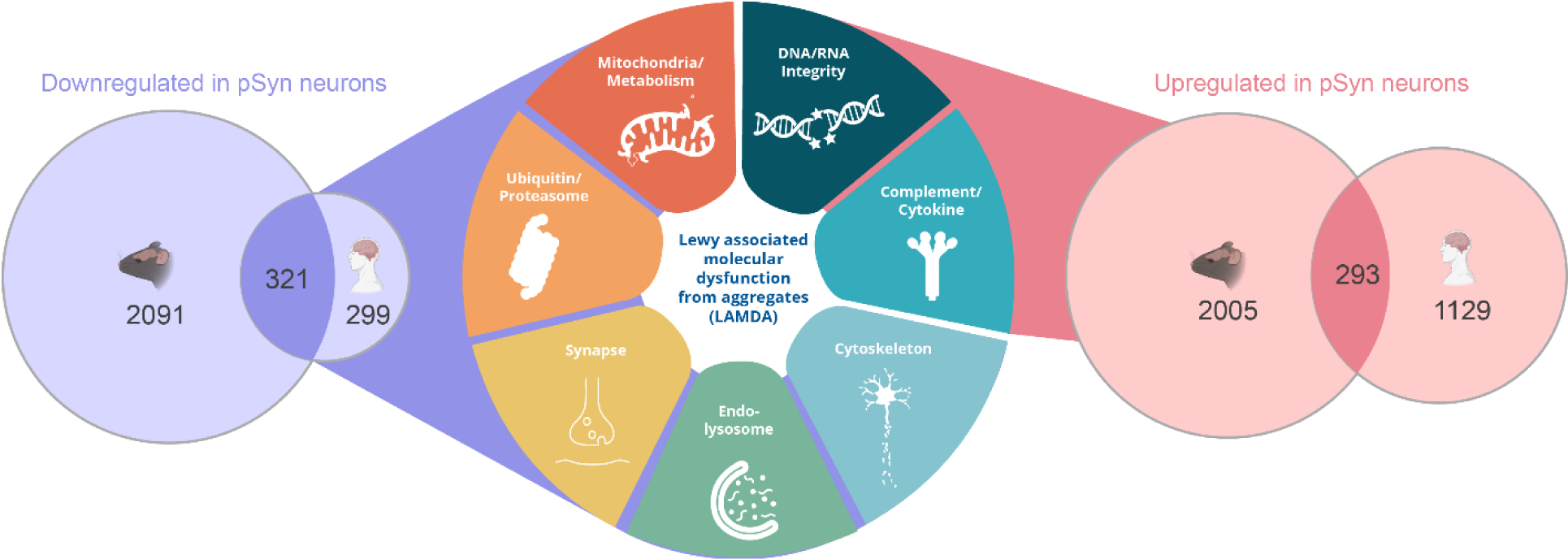
Lewy-associated molecular dysfunction from aggregates (LAMDA) Numbers of genes downregulated or upregulated in pSyn inclusion-bearing neurons are plotted from mouse and human tissue. The overlapping area is representative of the genes which show conserved expression changes in mice and humans. Conserved upregulated and downregulated genes in mouse and human fall with certain pathways and are described as a Lewy associated molecular dysfunction from aggregates (LAMDA) signature.

